# Re-coding of G protein-coupled receptor signaling enables emergent cellular behavior

**DOI:** 10.1101/2023.12.18.572241

**Authors:** Farhad Dehkhoda, Mitchell T. Ringuet, Emily A. Whitfield, Keith Mutunduwe, Cameron J. Nowell, Desye Misganaw, Zheng Xu, Akhter M. Hossain, Linda J. Fothergill, Stuart J. McDougall, John B. Furness, Sebastian G.B. Furness

## Abstract

All cells face the challenge of integrating multiple extracellular signals to produce relevant physiological responses. Different combinations of G protein-coupled receptors, when co-expressed, can lead to distinct cellular outputs, yet the molecular basis for this co-operativity is controversial. One such interaction is the reversal, from inhibition to excitation, at the dopamine D2 receptor in the ghrelin receptor’s presence, relevant for defecation control. Here we demonstrate that this reversal of dopamine D2 activity, to excitatory, occurs through a dominant switch in downstream signaling. This dominant switch, mediated by downstream signaling, enables fidelity in cellular responses not possible under alternative models, and provides an explanation for previously unresolved observations. Importantly, the switch in D2 signaling does not require ghrelin receptor agonism, rather its constitutive activity, thus accounting for the importance of central nervous system-ghrelin receptor in the absence of endogenous ligands. This re-coding has important implications for our understanding of how atypical receptor pharmacology can occur as well as how sequential signaling at individual neurons may be encoded to produce new outputs.

## Introduction

How cells interpret the multitude of cues they receive from the extracellular environment to produce the appropriate physiological response is still being elucidated. Much of neuronal signaling requires the coordination of outputs from multiple G protein-coupled receptors (GPCRs) and ion channels. This presents an immediate challenge to neurons because, while the vast majority of GPCRs perform non-overlapping physiological functions, a typical neuron expresses about 80 different GPCRs^1,2^, yet these couple to a limited palette of primary transducers^3^. At the molecular level, how the activity of different GPCRs is coded to produce distinct cellular, and thus physiological, outputs is contentious ^4,5^. The Rhodopsin-like Class A are the largest GPCR sub-family^6^, with the majority binding their ligands in the transmembrane bundle and exhibiting a common activation mechanism^7^. While class A GPCRs do not have an absolute requirement for dimerization for function^8–10^, there is a substantial amount of evidence that they can form dimers and higher order oligomers^11^. This has been proposed as a mechanism to explain atypical receptor pharmacology and physiology. Some have cast doubt on this model, highlighting that most studies on GPCR dimerization rely on overexpression with reported heterodimers a possible product of receptor crowding and random collisions^4,5^, but there is little data to support alternatives. We^12,13^ and others^14^, have shown that a subset of neurons that use dopamine as a neuromodulator at the dopamine D2 receptor (DRD2) also express the ghrelin receptor (GHSR), both Rhodopsin-like GPCRs. At these GHSR expressing neurons, dopamine displays an atypical excitatory response^15^. The DRD2 is typically coupled to Gαi/o^16,17^ isoforms, and in most neurons DRD2 activation promotes release of Gαi/o associated Gβγ subunits which open G protein-coupled inwardly rectifying potassium (GIRK) channels leading to membrane hyperpolarization and reduced excitability^18^. GHSR, on the other hand, is best characterized as coupling to Gαq/11 isoforms, which, when activated stimulate phospholipase C (PLC) activity^19^. In neurons, the coordinated effect of phosphatidylinositol 4,5-bisphosphate depletion and release of calcium from intracellular stores (via inositol-3-phophate (IP3) leads to M-type potassium channel closure and subsequent membrane depolarization^20^. Thus, in GHSR expressing neurons of the lumbrosacral defecation center, dopamine’s excitatory activity at DRD2 is more typical of Gαq/11 coupling, rather than Gαi/o coupling, where Gαi/o coupling is thought to be the predominant signaling pathway from DRD2^17^. We therefore sought to characterize, at the cellular and mechanistic level, how and why DRD2 signaling in the presence of GHSR is re-coded, which is informative of atypical receptor physiology more broadly.

## Results

### Dopamine-induced excitation of DRD2 on mouse lumbosacral spinal cord neurons is reversed with the depletion of intracellular calcium

Previous work by our group has shown that neurons in the spinal defecation centre of both rats and mice that are excited by dopamine are also excited by GHSR agonism ^13^ and that agonists acting on spinal defecation centers are capable of eliciting colorectal contractile activity ^12,21,22^. While DRD2 appears to be a physiologically relevant regulator for co-ordination of voluntary defecation in the spinal defecation center ^15^, GHSR antagonism can oppose this effect ^12,21^, which is supported by recent work of Sawamura et. al. ^23^. We have also shown that, in spinal defecation center neurons in neonatal rats and adult mice, ∼85% respond to dopamine with ∼25% exhibiting typical inhibitory responses and ∼60% an atypical excitatory response ^13^. In defecation center neurons from the same animals ∼75% are excited by GHSR activation by capromorelin ^13^ and the excitatory effect of dopamine is blocked by GHSR antagonism ^13^. Based on previous reports ^14,24^ and the accepted mechanism of GPCR-dependent neuronal depolarization (e.g. ^25^), we hypothesized that intracellular Ca^2+^ mobilization (_i_[Ca^2+^]) in *Drd2* neurons in the spinal defecation center was at least in part responsible for the excitation we observed. We identified *Drd2* positive neurons in the spinal defecation center using a drd2-tdTomato reporter mouse and incubated coronal slices in Calbryte 520AM for _i_[Ca^2+^] imaging (Fig. 1A). Our results revealed that _i_[Ca^2+^] mobilization occurred in *Drd2* neurons following dopamine (30 μM) and KCl (75 mM) wash on (Fig. 1B, C). _i_[Ca^2+^] store depletion using thapsigargin reversed the excitatory effect of dopamine at DRD2 and reduced capromorelin’s excitatory effect (Fig. 1D, E), compared to the expected response of dopamine and capromorelin’s excitatory effect in the same neuron (Fig. 1D, E). Thus, DRD2 activation still produced an electrophysiologically measurable, but reversed, output, whereas GHSR effects were reduced. These findings indicate that DRD2 and GHSR have parallel excitatory effects when store Ca^2+^ is available; and that in the absence of store Ca^2+^ these receptors can operate independently in neurons of the lumbosacral defecation centers.

**Fig. 1.**
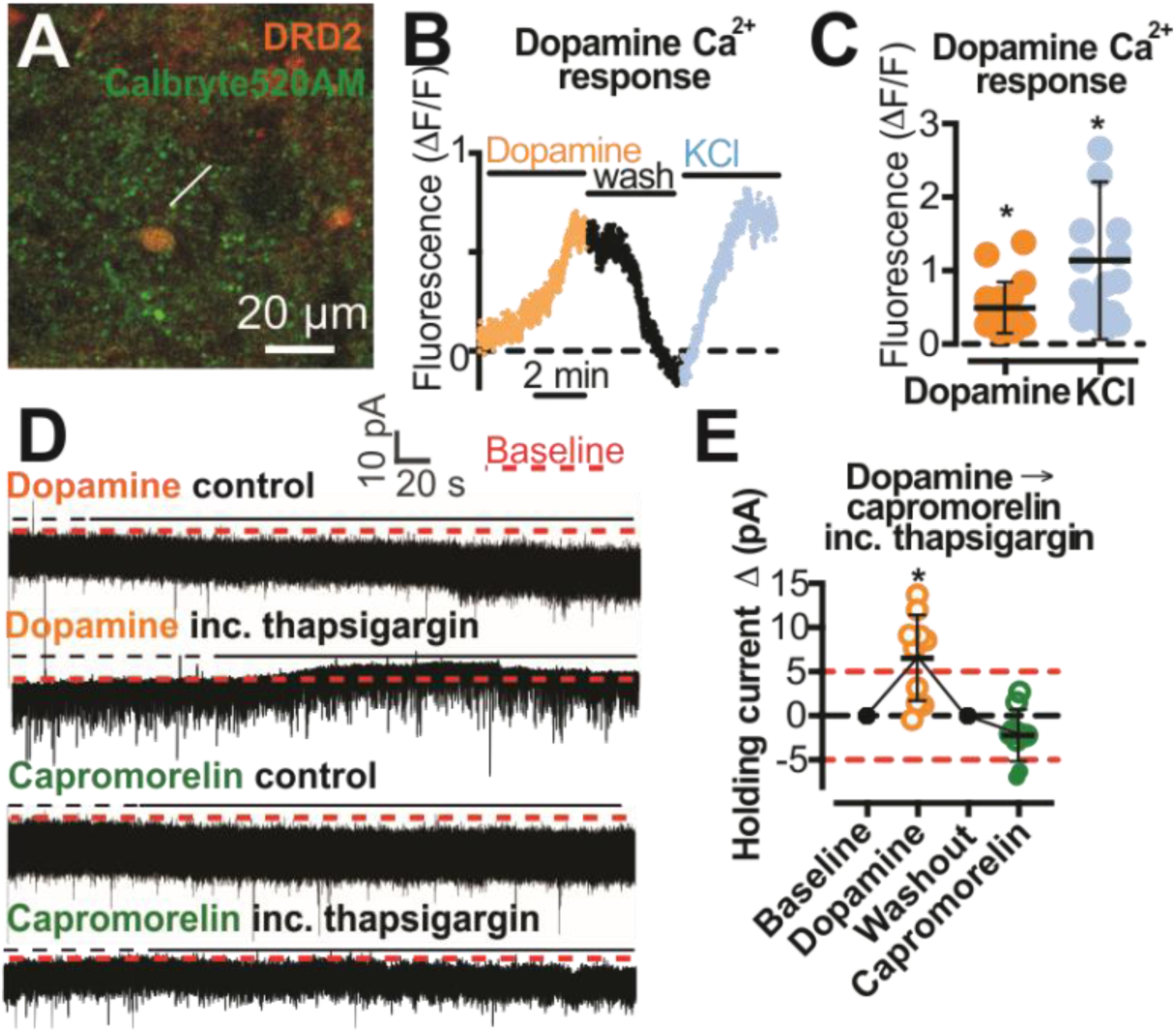
Activation of DRD2 or GHSR on spinal defecation center neurons is differentially affected by calcium store depletion. (**A**) Drd2-tdTomato-positive neuron (DRD2; orange) filled with the synthetic calcium indicator, Calbryte 520AM (5 μM; green).(**B**) Representative fluorescence trace of DRD2 neuron showing Ca^2+^ responses following dopamine (30 μΜ; orange) and KCl (75 mM). (**C)** Peak Ca^2+^ fluorescence following dopamine (30 μM) and KCl (75 mM), n = 4 mice, 18 neurons. (**D**) Representative DRD2 neuron recordings show inward (excitatory) current in response to dopamine (30 μΜ) in control and outward current (inhibition) in the presence of thapsigargin (1 μM). The inward current response in response to 10 nM capromorelin was not reversed in the presence of 1 μΜ thapsigargin. (**E**) Dopamine (30 μM; orange) generates outward currents, while capromorelin (10 nM; green) exhibited reduced inward currents in the presence of thapsigargin (1 μM) in the same DRD2 neurons. Red dashed lines at −5 pA and 5 pA are deemed excitatory and inhibitory thresholds. All data are mean ± SD, paired student’s t-test, peak of drug response delta holding I (pA) /washout, * p<0.05.

### GHSR leads to potent re-coding of cellular responses to DRD2 activation in recombinant cell lines

Observations in native cells indicate that DRD2 activity is re-coded from an inhibitory to an excitatory output. Our previous data indicate that this re-coding was blocked by GHSR antagonism ^13^ and the present data indicates it requires _i_[Ca^2+^] mobilization (Fig. 1D, E). Previous studies have suggested that re-coding of DRD2 responses result from a dimerization-dependent switch in primary transducer ^14^.. We therefore next investigated this in controlled stable cell lines, in which genes for either receptor were present as single copies, resulting in moderate, rather than high receptor expression (Fig. 2A, B, supp Fig. 2J, K). The functional interaction that we had observed in spinal neurons was preserved in transfected cells, with atypical DRD2 signaling occurring due to the presence of GHSR (Fig. 2C-D, Supp 2A, L). In the absence of GHSR, dopamine (Fig. 2C, D, G) elicited a mean calcium peak of 3.8% of ionomycin, which increased to a mean of 32% in the presence of GHSR (logEC_50_ −8.3, ∼5 nM, Fig. 2D, Supp 2C). The potency of dopamine to elicit a calcium response was very close to that for inhibition of forskolin mediated cAMP (∼11nM, Fig. 2J), suggesting a similar ability to engage the transducer(s) responsible for each pathway. A similar effect was obtained with the DRD2-preferring agonist pramipexole (supp Fig 2A, D, E, H, I), and we observed that the presence of either receptor does not alter the cell surface expression of the other (Fig. 2A-B, Supp 2J, K). The magnitudes and kinetic differences of calcium release from DRD2 and GHSR activation were consistent with different pathways to _i_[Ca^2+^] mobilization (Fig. 2C-F). Although _i_[Ca^2+^] downstream of DRD2 was GHSR dependent (Fig. 2C-D), cAMP inhibition was not, shown by the presence of GHSR having no statistically significant or biologically relevant effect on the ability of DRD2 ligands to inhibit the activity of forskolin-stimulated adenylate cyclase (Fig. 2I, J, Supp 2H, I). These results suggest that GHSR provides a dominant second messenger switch for DRD2 in recombinant cell lines (as it does in native cells), consistent with a switch downstream of DRD2’s primary transducer.

**Fig. 2.**
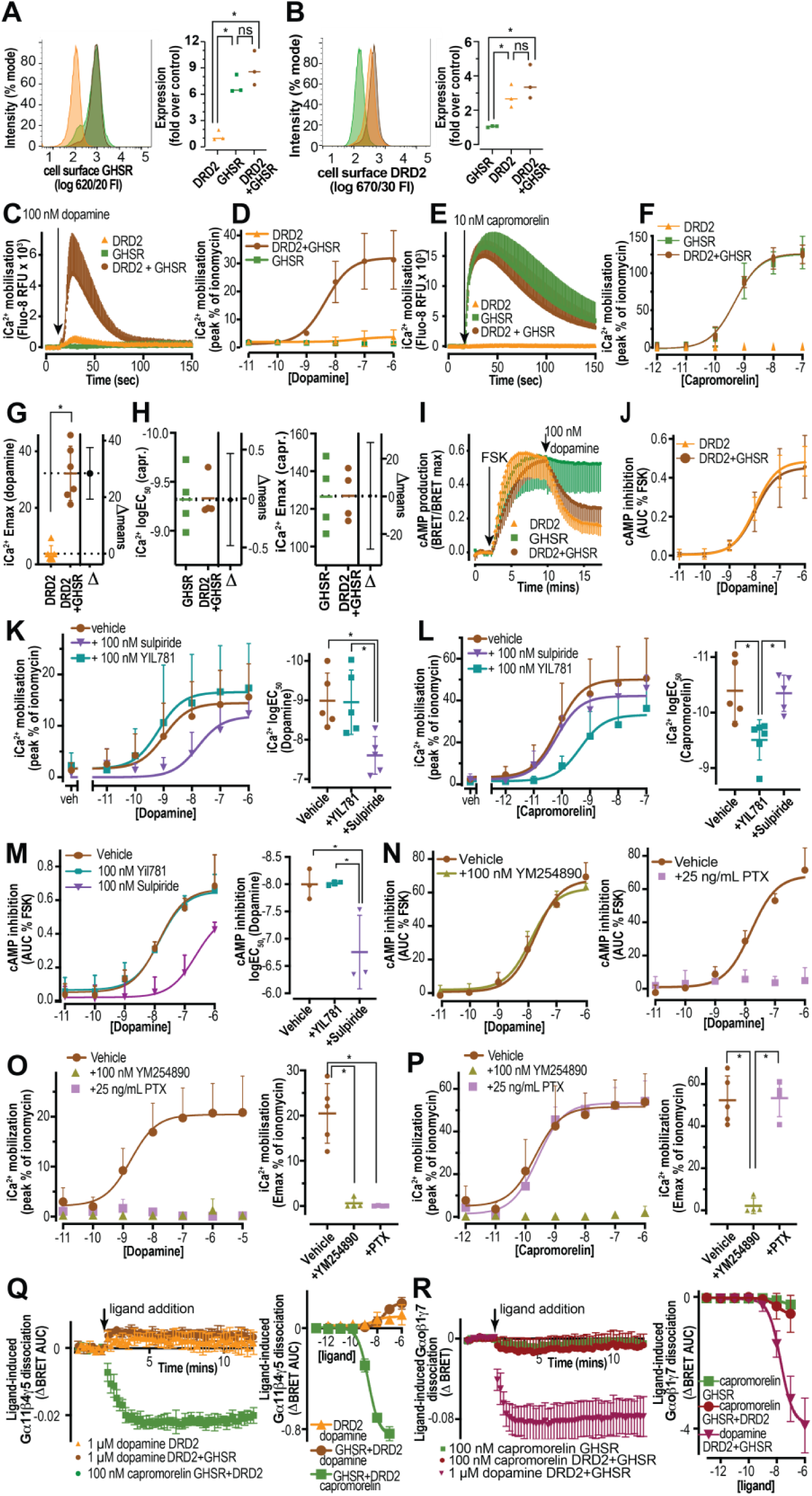
GHSR provides a pharmacological switch for DRD2 in recombinant cell lines but pathways from GHSR and DRD2 are unique. (**A,B**) Cell surface expression is not altered by co-expression. Cell surface expression of GHSR (**A**) and DRD2 (**B**) in single copy stably transfected flpIN CHO cell lines using flow cytometry and pooled (fold over control), respectively, in DRD2 (orange, n=3), GHSR (green, n=3) and DRD2+GHSR (brown, n=3). (**C**) GHSR expression is permissive for intracellular _i_[Ca^2+^] response of DRD2. Calcium increase (raw RFU) is seen following application of 100 nM dopamine in DRD2+GHSR-transfected cells (brown), but is minimal in DRD2 alone cells (orange) and undetectable in GHSR alone cells (green) alone (n=5). (**D**) _i_[Ca^2+^] quantified as a peak % of ionomycin following dopamine application in DRD2 (orange, n=5), GHSR (green, n=5), and DRD2+GHSR (brown, n=5) transfected cells. (**E**) DRD2 does not change coupling of GHSR to calcium. Ca^2+^ increase following application of 10 nM capromorelin in DRD2+GHSR (brown) and GHSR (green) but is absent in DRD2 (orange) transfected cells (n=5). (**F**) _i_[Ca^2+^] quantified as a peak % of ionomycin following capromorelin application in DRD2 (orange, n=5), GHSR (green, n=5), and DRD2 + GHSR (brown, n=5) transfected cells. (**G,H**) Quantification of EC_50_ and E_max_ of _i_[Ca^2+^] in DRD2 (orange) and DRD2+GHSR (brown) transfected cells following dopamine (**G**) or capromorelin (Capr., **H**) application, based on data in (**D and E**). (**I**) GHSR does not alter DRD2 coupling to cAMP inhibition. Forskolin (FSK; 3 µM) mediated increase of cAMP (BRET/BRET max) which is reduced by dopamine (100 nM) in cells transfected with DRD2+GHSR (brown) or with DRD2 (orange), n=5 but not GHSR alone. (**J**) quantification using area under the curve (AUC) % of FSK as concentration response to dopamine. **(K, L)** The effect of DRD2 preferring antagonist, sulpiride and GHSR antagonist, YIL781 on dopamine (**K**) and capromorelin (**L**) dependent _i_[Ca^2+^], concentration-response curves of peak % ionomycin response with quantified EC_50_ (n=5). **(M)** Reduction with DRD2 preferring antagonist, sulpiride and no effect of GHSR antagonist, YIL781 on dopamine-dependent inhibition of cAMP production, concentration-response curves AUC % inhibition of FSK response with quantified EC_50_ with (n=5). (**N**) The effect of Ga_q/11_ (YM254890, left panel) or Ga_i/o_ (PTX, pertussis toxin) inhibition on dopamine-dependent inhibition of cAMP production, concentration-response curves AUC % inhibition of FSK response (n=3). (**O, P**) The effect of Ga_q/11_ (YM254890) or Ga_i/o_ (PTX) inhibition on dopamine- (**O**) or capromorelin- (**P**) dependent _i_[Ca^2+^], as concentration-response curves peak % of ionomycin response with quantified E_max_ (n=5). (**Q**) The presence of GHSR is not permissive for DRD2 coupling to Ga_11_/β4γ5 using TRUPATH G protein dissociation assay, left kinetic traces of BRET change in response to saturating ligand and right concentration response curves. (**R**) The presence of DRD2 is not permissive for GHSR coupling to Ga_o_/β1γ7 using a TRUPATH G protein dissociation assay, left kinetic traces of BRET change in response to saturating ligand and right concentration response curves. All data are mean ± SD or mean +SD; * p<0.05; ns, not significant, one-way ANOVA or unpaired t-test, with estimation plot.

### GHSR and DRD2 have independent signaling pathways in recombinant cell lines

Because our data on second messenger signaling is consistent with unchanged DRD2 primary transducer coupling in the presence of GHSR, we sought to meticulously characterize this coupling. We began by examining the nature of the re-coding at a receptor-level. In cells containing GHSR and DRD2, antagonism of DRD2 blocks dopamine induced _i_[Ca^2+^] mobilization (Fig. 2K) and reduces cAMP inhibition (Fig. 2M), producing comparable 1.38 and 1.24 log reductions in mean EC_50_s, but has no effect on GHSR _i_[Ca^2+^] mobilization (capromorelin stimulation, Fig. 2J).

Comparable data was seen for pramipexole and ghrelin (Supp Fig. 3A, B, E). Furthermore, GHSR blockade with YIL-781 does not affect DRD2 calcium coupling (Fig. 2K) or cAMP inhibition (Fig. 2M) and produces an expected 0.9 log right-shift in _i_[Ca^2+^] mobilization induced by capromorelin (Fig. 2L), which was comparable for ghrelin (Supp Fig 3C, D). While YIL-781 is reported to be a GHSR inverse agonist, Mende et al. ^26^ report this to be a weak partial agonist for Gαq/11 in highly coupled systems. Taken together these data are consistent with a dominant second messenger switch of DRD2 provided by GHSR rather than a change in primary transducer via receptor dimerization. Therefore, we sought to understand the involvement of Gαi/o, DRD2’s typical coupling partner, and Gαq/11, the typical partner for GHSR, in _i_[Ca^2+^] mobilization. Inhibition of either Gαi/o with pertussis toxin (PTX) or Gαq/11 with YM254890 ablated dopamine (Fig 2O) and pramipexole (Supp Fig 3I, J) dependent _i_[Ca^2+^] mobilization in the GHSR plus DRD2 expressing cell-line, indicating that *both* Gαi/o and Gαq/11 are essential for DRD2-dependent _i_[Ca^2+^] mobilization. By contrast Gαq/11 inhibition with YM254890 had no detectable effect on dopamine (Fig 2N) or pramipexole (Supp Fig 3G) induced inhibition of cAMP production, while, as expected, this cAMP response was completely ablated by PTX (Fig 2N and Supp Fig 3H). Capromorelin (Fig 2P) and ghrelin (Supp Fig 3K, L) dependent _i_[Ca^2+^] mobilization was unaffected by inhibition of Gαi/o with PTX but was undetectable when cells were pre-treated with YM254890. We also used the TRUPATH system ^27^ to measure whether we could detect a change in preference for coupling to these exogenous G protein sensors. Although DRD2 coupled well to Gαo (Fig 1R, Supp Fig 3O, P) and GHSR to Gα11 (Fig 1Q, Supp Fig 3M, N), our results revealed that the co-expression of both receptors was not permissive for either receptor to couple to its non-preferred partner. Together these data are consistent with there being no change in the primary coupling of either receptor to their preferred Gα partner in the presence of the other receptor and with a co-operative effect of Gαi/o activation (downstream of DRD2) with basal Gαq/11 activity.

**Fig. 3.**
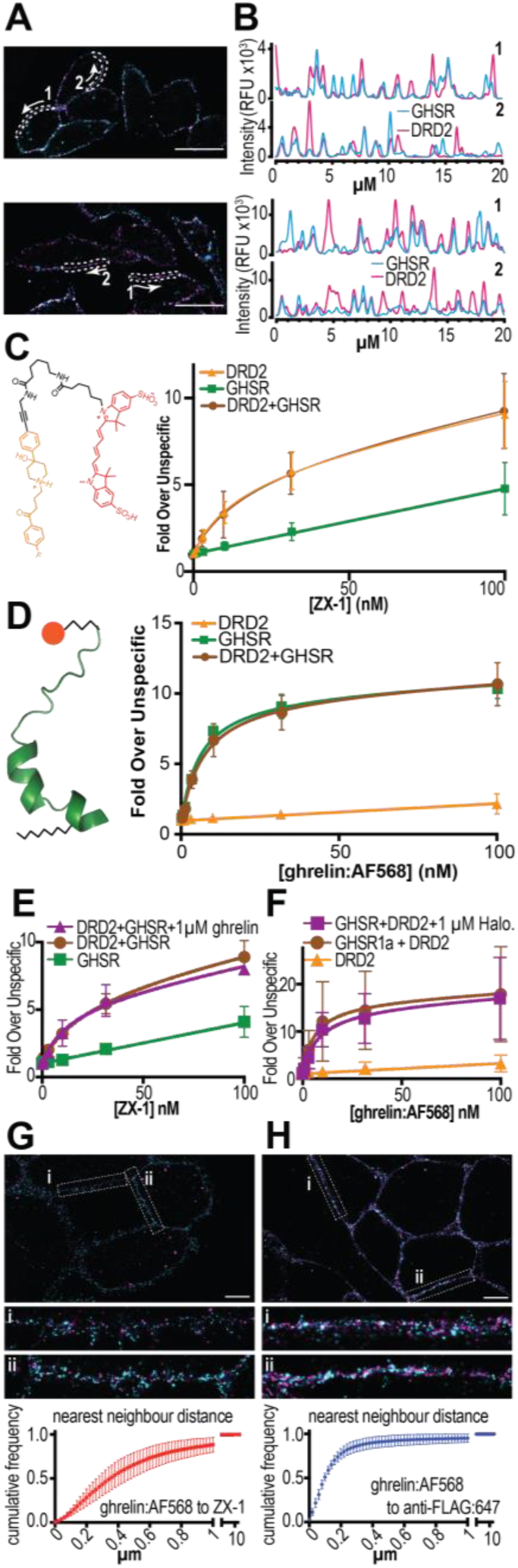
GHSR and DRD2 appear to be independent at the cell surface in recombinant cell lines. (**A**) Representative deconvolution microscopy (n=3) using direct conjugate antibodies directed toward n-terminal tags of DRD2 (pseudo-colored in magenta) and GHSR (pseudo-colored in cyan) with effective resolution, xy=160 nm, z=385 nm, scale bars are 20 µM. (**B**) Pixel intensity plots for each channel in the selected membrane segments indicated in (**A**), in the directions of the arrows. (**C**) Structure of ZX-1, a novel DRD2-preferring fluorescent ligand comprising haloperidol (orange) linked (black) to sulfo-Cy5 (red) and its saturation ligand biding by flow cytometry (no-wash protocol) (n=7). (**D**) Saturation ligand biding by flow cytometry (no-wash protocol) using ghrelin:AF568, (n=5). Ghrelin:AF568 model at left. (**E**) Saturation binding by flow cytometry (no-wash protocol) using ZX-1 in the absence and presence of excess (1µM) ghrelin (n=3). (**F**) Saturation binding by flow cytometry (no-wash protocol) using ghrelin:AF568 in the absence and presence of excess (1µM) haloperidol (n=3). (**G, H**) Representative super-resolution image using fluorescently labeled DRD2 and GHSR ligand staining (**G**) and anti-FLAG and GHSR ligand staining (**H**) of DRD2+GHSR (**G**) and GHSR (**H**) cells with an effective resolution xy=15 nm z=42 nm. **i** and **ii** show a zoom of areas of the presumptive cell membrane. Bottom shows analysis of pixel-by-pixel minimum distances to nearest neighbor in alternative channel (n=5 independent staining/imaging each with 2-3 images). Scale bar is 5 µM, ghrelin:AF568 is pseudo-colored in cyan, ZX-1 and anti-FLAG:AF647 are pseudo-colored in magenta. All data are mean ± SD.

### No evidence for GHSR and DRD2 direct interaction in recombinant cell lines

Our interpretation of the reversal of neuronal activity in the presence of GHSR blockade, and the molecular pharmacological analysis, is of a second messenger interaction downstream of DRD2 and GHSR. This dominant switch in second messenger coupling could, nonetheless, be attributed to receptor dimerization. Using directly conjugated antibodies, we performed qualitative deconvolution microscopy to examine the distributions of GHSR and DRD2 (Fig 3A, B, Supp Fig 4 A-C). Although we observed a degree of overlapping signal, for the most part this staining methodology did not support a general co-localization of the two receptors. As an alternative approach, we performed live-cell saturation binding to measure an allosteric effect of one receptor on the other. Both DRD2 and GHSR are essentially transmembrane bundle only receptors, with ligand binding pockets in the transmembrane bundle. A dimerization-dependent switch in DRD2 primary coupling would be expected to involve transmembrane domain allostery with ligand binding co-operativity viewed as the strongest evidence for heterodimers ^28^. Based on the structure of DRD2 bound to haloperidol (6LUQ^29^) we reasoned that the haloperidol core could be extended to allow fluorophore conjugation (sulfo-Cy5) to generate a DRD2-preferring fluorescent ligand (shown in Fig 3C, left panel, orange:haloperidol, black:linker, red:sulfo-Cy5, compound name – ZX-1, Supp Fig. 4D). We then used this compound to perform saturation binding on cells expressing DRD2 alone or DRD2 plus GHSR (Fig 3C). No difference in DRD2 affinity for this ligand in the absence or presence of GHSR (Fig 3C, Supp Fig 4E, F) was detected and the binding data reported the same DRD2 receptor number (Fig 3C, Supp Fig 4E, F), independent of GHSR expression, consistent with our flow cytometry results (Fig 2B). Similarly, our published fluorescent GHSR ligand (ghrelin:AF568^30^) could not detect any effect of DRD2 expression on either the affinity of GHSR for this ligand or the number of available GHSRs at the cell surface (Fig 3D, Supp Fig. 4G, H, consistent with Fig 2A). Moreover, saturating concentrations of unlabeled haloperidol had no detectable effect on the binding of ghrelin:AF568 to GHSR (Fig 3E, Sup Fig. 4J) and nor did unlabeled ghrelin affect the binding of ZX-1 to DRD2 (Fig 3F, Supp Fig. 4I), indicating no functionally detectable allosteric interaction between DRD2 and GHSR in these single-gene-copy cells. As these fluorescent ligands provide chemically defined, singly labeled tools where the fluorophore will be very close to the receptor binding pocket when bound, we reasoned that these might be suitable for fine-grained analysis of the distribution of GHSR and DRD2 by super-resolution microscopy. We performed staining and fixation of these cells prior to imaging using STochastic Optical Reconstruction Microscopy (STORM) which were processed, drift-corrected and channel aligned to yield images (vector maps) with effective xy resolutions of 15 nm (Fig 3G). Visual inspection suggested almost none of the localized ZX-1 and ghrelin:AF568 were within the distance range expected for DRD2:GHSR dimerization (Fig 3G). We performed analysis using direct measurement of nearest neighbor distance of ghrelin:AF568 to ZX-1 (3G, bottom panel), where ∼ 2% of all measurements were less than our generous cut-off for dimerization of 25 nm (3x receptor diameter plus allowance for ligand flexibility). This supports the notion that DRD2 and GHSR do not undergo significant heteromerization in these single-copy cells, however at higher receptor densities may place receptors near enough to allow resonance energy transfer using standard fusion protein or antibody methods providing a potential explanation for the discrepancy between our observation and those of others. As a control, Fig 3H shows the localization of ghrelin:AF568 compared to a direct conjugate anti-FLAG:AF647, directed toward the N-terminal tag of the ghrelin receptor. Visual inspection suggested some overlap of signal.

**Fig. 4.**
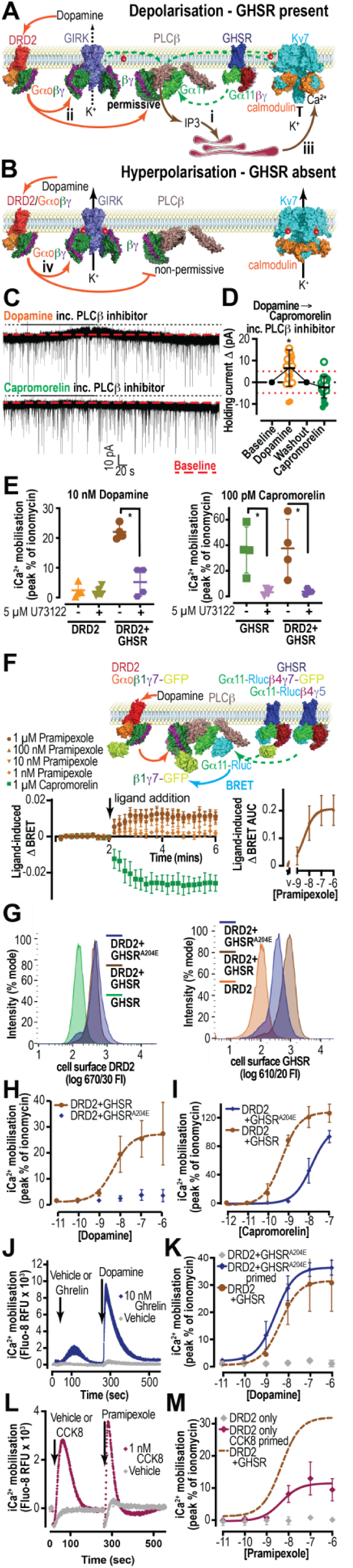
Model for DRD2 calcium coupling and its testing in neurons in the lumbosacral spinal cord, and the requirement for PLCβ. **(A)** Model for the GHSR-dependent switch in DRD2 activity, (**i**) the constitutive activity of GHSR coupled to Gαq/11 primes the system by partly depleting PIP2 from PLCβ, permissive to activation by Gαi/o-associated βγ subunits liberated by DRD2 activation (**ii**) which then fully activates PLCβ, leading to intracellular calcium mobilization and (**iii**) causing a conformational switch in calmodulin at Kv7 (M type) K^+^ channels, resulting in channel closure and membrane depolarization. (**B**) In the absence of GHSR-dependent PLCβ priming, DRD2 activation leads to typical Gαi/o associated βγ subunit activation of GIRK channels. (**C**) Representative voltage clamp trace from a PPN generating outward current (inhibition), relative to baseline holding current (red-dashed line, all experiments) following 30 μΜ dopamine (top) and minimal effect of 10 nM capromorelin (bottom) wash on in the presence of PLCβ inhibitor (10 μM; U73122). Solid line indicates deduced drug effect period. (**D**) Dopamine (30 μΜ) and capromorelin (10 nM) successive application in the presence of U73122 (10 μM). In most neurons, dopamine caused an outward or no current (open circles). Closed circles indicate neurons where the inward current was >-5 pA. Capromorelin caused no effect or a small outward current in most neurons (open circles) or inward current (closed circles), that was on average smaller than control (Fig 1E). Red dashed line at −5 pA is deemed the excitatory threshold. Red dashed line at 5 pA is deemed the inhibitory threshold. (**E**) In transfected cells, co-addition of 5µM U73122 (PLCβ inhibitor) largely blocks the ability of dopamine and capromorelin to evoke intracellular Ca^2+^ mobilization, shown as scatter plot of peak calcium (n=5). (**F**) In a TRUPATH assay using Ga_11Rluc_/β4γ5 in combination with Ga_o_/β1γ7_GFP_, activation of DRD2 with pramipexole results in a BRET increase, while capromorelin stimulation of GHSR causes a decrease (model at top, kinetics at left, concentration-response AUC at right (n=4). (**G**) GHSR^A204E^ (mutant with no constitutive activity) does not alter cell surface expression of DRD2 but is expressed at a lower level compared with wild-type GHSR; cell surface expression of DRD2 and GHSR^A204E^ in single copy stably transfected cell lines using flow cytometry, left expression of DRD2 and right expression of GHSR, in GHSR (green, n=3), GHSR+DRD2 (brown, n=3) and DRD2+ GHSR^A204E^ (purple, n=3) cell lines. (**H**) DRD2 fails to couple to Ca^2+^ in the presence of GHSR^A204E^ following dopamine stimulation (**I**) and capromorelin-induced calcium coupling is right shifted for GHSR^A204E^ compared to conventional GHSR. Mean ± SD. (n=3). (**J**) DRD2 Ca^2+^ coupling is rescued by pre-stimulation of GHSR^A204E^ with low concentration ghrelin. Representative trace showing pre-stimulation with ghrelin (10 nM; blue, n=3) eliciting a small Ca^2+^ response compared to vehicle (gray), followed by dopamine leading to a sharp rise and substantial increase in Ca^2+^ release, compared to vehicle (gray). (**K**) Dose-response relationship of Ca^2+^ increase from dopamine following pre-stimulation with ghrelin (10nM; blue, n=3). (**L**) DRD2 Ca^2+^ coupling in DRD2 only cells is rescued by pre-stimulation of endogenous CCK1-R with low concentration CCK-8. Representative trace showing pre-stimulation with CCK-8 (1 nM; magenta, n=3) eliciting a Ca^2+^ response compared to vehicle (gray), followed by pramipexole leading to a sharp rise and substantial increase in Ca^2+^ release, compared to vehicle (gray). (**M**) Dose-response relationship of Ca^2+^ increase from pramipexole following pre-stimulation with CCK-8 (1 nM; magenta, n=3). Mean ± SD, paired student’s t-test (**D**), one way ANOVA (**E**) * p<0.05

Direct nearest neighbor measurements show >30% of all measurements were within 60 nm (1x receptor plus antibody size plus unstructured n-terminus of GHSR plus allowance for ghrelin:AF568 flexibility) for co-binding of ghrelin:AF568 and anti-FLAG:647 to the same receptor. The difference in proximity between fluorescently labelled ligands and a fluorescently labelled ligand and antibody are illustrated in Supp Fig 4K.

### DRD2 calcium coupling in neurons in the lumbosacral spinal cord requires PLCβ

A common mechanism whereby GPCRs couple to neuronal depolarization is typified by the ability of Gαq/11 to activate PLCβ which in turn inactivates Kv7 (M-type) potassium channels ^25^. Based on the above data, this known mechanism for GPCR-dependent depolarization, and the recent data from the Kostenis’ lab ^31^, demonstrating the requirement for Gαq/11 priming to allow PLCβ to be susceptible to activation by Gαi/o-associated βγ, we propose the model shown in Figure 4A & B. In this model, dopamine causes depolarization due to GHSR-dependent Gαq/11 priming of PLCβ, which enables PLCβ activation by βγ released from DRD2, dominantly inhibiting M-type potassium channels. Contrastingly, in the absence of PLCβ priming and M-channel inhibition, βγ released from DRD2 can dominantly activate GIRK channels, leading to hyperpolarization. Thus, blockade of GHSR and depletion of calcium stores would be expected to lead to DRD2-dependent hyperpolarization (Fig 1 and ^13^), the DRD2-dependent increase in iCa^2+^ would be dependent on the presence of GHSR (Fig 2) as well as both Gαi/o and Gαq/11 (Fig 2) but cAMP coupling via Gαi/o would be unaffected by the presence of GHSR (Fig 2) and there would be no requirement for direct receptor:receptor interaction (Fig 3). This model implies that blockade of PLCβ should also lead to DRD2-dependent hyperpolarization and indeed, we observed that in the presence of a PLCβ inhibitor, U73122, dopamine failed to elicit atypical inward excitatory currents, and instead caused typical, outward inhibitory currents (Fig. 4C, D). Consistent with our model, PLCβ inhibition also reduced capromorelin-induced inward, excitatory currents (Fig. 4C, D) an effect that was observed in the same PPN.

### GHSR-dependent coupling of DRD2 to calcium requires GHSR constitutive activity

We sought to confirm integration of DRD2 and GHSR signaling at PLCβ. Consistent with neuronal activity, the PLCβ inhibitor, U73122, blocked both DRD2- and GHSR-dependent Ca^2+^ mobilization in transfected cells (Fig 4G and Supp Fig 5A). Furthermore, when we combined TRUPATH sensors for both Gαo and Gα11, but with the donor Rluc8 only on Gα11 and the acceptor GFP on the Gαo preferred Gγ7, stimulation of DRD2 in cells co-expressing GHSR caused an increase in BRET signal, consistent with an increase in proximity of Gα11_Rluc8_ and Gγ7_GFP_, (Fig 4H). This is consistent with our model where Gα11 and βγ subunits liberated from Gαo converge on PLCβ (Fig 4A and H). Because this integration of signaling and our model requires GHSR constitutive activity, we developed single-gene copy cell lines that co-express DRD2 and the naturally occurring GHSR^A204E^ mutant, which is reported to lack constitutive activity, but is still responsive to ghrelin ^32^. This mutant did not alter cell surface expression of DRD2 (Fig 4H, left panel, quantitated Supp 5B) but exhibited 2-fold lower expression compared with the wild-type receptor (Fig 4H, quantitated supp 5B). Co-expression of GHSR^A204E^ with DRD was completely non-permissive for DRD2-dependent calcium mobilization (Fig 4I and Supp Fig 5C), and also displayed reduced responsiveness to capromorelin (Fig 4J) and ghrelin (Supp Fig 5D). Conversely, system priming with an approximate EC_10_ concentration of ghrelin allowed dopamine (Fig 4K, L) and pramipexole (Supp Fig 5E, F) to couple to intracellular calcium mobilization to the same extent as in cells co-expressing wild-type GHSR, supporting the notion that the low-level tonic activation of Gαq/11, downstream of GHSR, is necessary and sufficient for a pharmacological switch in DRD2 behavior. This suggested to us that system priming via another Gαq/11 coupled GPCR may be permissive for DRD2-dependent Ca^2+^ mobilization. These cells express low, but pharmacologically detectable levels of the cholecyctokinin 1 receptor (CCK1R), a typically Gαq/11 coupled GPCR. Priming of DRD2-only expressing cells with a low (1 nM, ∼EC_30_) concentration of CCK-8 led to a recovery in the ability of dopamine to elicit a Ca^2+^ response, with an EC_50_ similar to that seen in cells co-expressing DRD2 with GHSR. This suggests a generalizable effect of Gαq/11 priming on the ability of DRD2 to couple to calcium via Gαi/o.

## Discussion

DRD2s are expressed in a subset of neurons throughout the central nervous system, where they participate in a range of different physiologies including motor control, reward (including feeding reward), cognition and, as we and others have recently shown, voluntary defecation ^15,23^. These receptors typically act by inhibiting neuronal excitation through coupling to Gαi/o and downstream βγ dependent opening of GIRK channels ^18,33^. It has been reported that when DRD2 is associated with the ghrelin receptor, activation of DRD2 causes atypical excitation ^14,15,24^. We investigated the molecular mechanism through which this reversal of effect at DRD2 occurs in both transfected cells and native autonomic neurons. Our data supports a model in which the GHSR-dependent switch of DRD2’s action from inhibition to excitation depends on the ability of GHSR to tonically activate Gαq/11, such that Gαq/11-activated PLCβ is permissive for DRD2-dependent Gαi/o associated βγ activation. In contrast to previous reports ^14,24^, we found no evidence for dimerization being associated with the GHSR-dependent switch in DRD2’s action. Our results also demonstrate that the re-coding of DRD2 can occur through pre-stimulation of the naturally occurring GHSR^A204E^ mutant that lacks constitutive activity, but is still agonist responsive, with a very low concentration of ghrelin. This enables potent and efficacious coupling of DRD2 to an atypical calcium response. This ability of pre-stimulation of a Gαq/11 coupled receptor to re-code DRD2 response is also true for CCK1R.

These two re-coding effects have broad implications for GPCR biology. Firstly, in the case of GHSR recoding, a variety of observations regarding the ability of GHSR inverse agonists to alter dopaminergic and opioid signaling may be due to the physiological function of GHSR which allows these Gαi/o coupled receptors to have a neuro-excitatory effect. This has important implications for the choice of centrally penetrant GHSR antagonist for treating addiction disorders ^34^. Our model may account for the previous observation of loss of effectiveness of GHSR agonism after lesion of dopaminergic neurons in a preclinical model of Parkinson’s Disease^35^. We would propose that Parkinson’s Disease-associated chronic-constipation (present in 70-80% of patients^36^) may respond to subtherapeutic doses of capromorelin, which would act to potentiate the normal dopamine-mediated voluntary-defecation response. Another implication of GHSR dependent recoding of DRD2 activity with no change in primary transducer, is that it provides a mechanism for neuronal response fidelity, distinguishing an excitatory input from a Gαi/o coupled receptor (e.g. DRD2 in a GHSR expressing neuron) from a Gαq/11 coupled receptor (e.g. a muscarinic M1, 3 or 5 input). This mechanism for a fidelity in cellular response, does not require dimerization. In this example, both inputs may be expected to cause transient depolarization but because DRD2 activation will also cause a decrease in cellular cAMP, its activation might be expected to reduce after-depolarization events and produce different medium-term changes in gene expression. Secondly, the fact that very low-level pre-stimulation of Gαq/11, through both GHSR^A204E^ and CCK1R, leads to temporary re-coding of the DRD2 response implies a mechanism by which the order of signals arriving at a neuron could be coded to provide a physiologically plastic conditional response. The ability of GHSR to interact with other GPCRs could have implications in their physiological activity with important consequences for learning and memory, metabolic disorders and issues relating to colonic motility. This type of second-messenger convergence provides a model to understand observations such as the melanocortin-4 receptor-dependent reversal of valence that is mediated via the dopaminergic reward pathway ^37^. Thus, the ability to re-code GPCR output provides a mechanism by which emergent behavior can arise from a limited palette of transducers.

## Acknowledgments

The authors thank Ben Capuano, Peter Scammells and Rob Lane for their supervision of ZX and Ben Capuano for assistance with preparing methods relating to the compound ZX-1. The authors would like to acknowledge that the present work was conducted on the traditional lands of the Turrbal and Wurundjeri peoples. We would like to thank Dominic Ng, Sean Millard, Wally Thomas, Scott Prosser and Rowan Tweedale for constructive critique of the manuscript. STORM data acquisition was performed at the Queensland Brain Institute’s Advanced Microscopy Facility using the SAFe 360; Abbelight, France.

## Funding

2021 NHMRC Ideas Grant; 2021/GNT2012657 (SGBF, LJF, JBF)

SGBF is an ARC Future Fellow (FT180100543)

## Author contributions

Conceptualization: SGBF, JBF

Methodology: FD, MTR, EAW, CJN, ZX, AMH, SGBF

Investigation: FD, MTR, EAW, KM, DM, JBF, SGBF

Visualization: MTR, FD, CJN, JBF, SGBF

Funding acquisition: SGBF, LJF, JBF

Project administration: SGBF, JBF Supervision: SGBF, JBF, SJM

Writing – original draft: SGBF, JBF, FD, MTY

Writing – review & editing: FD, MTR, SGBF, JBF

## Data and materials availability

All data are available in the main text or the supplementary materials.

## Supplementary Materials

## Materials and Methods

### Animals

Experiments were conducted in accordance with the National Health and Medical Research Council guidelines for the care and use of animals and with approval from the Florey Institute of Neuroscience and Mental Health Animal Ethics Committee (FINMH-20-024). For whole-cell electrophysiology and calcium imaging experiments, adult D2R reporter mice (female and male; 8-20 weeks) were used. D2R-tdTomato (D2R) reporter mice were generated by crossing heterozygous Drd2-cre (B6.FVB(Cg)-Tg (Drd2-cre)ER44Gsat/Mmucd) with homozygous tdTomato (Ai14(RCL-tdT)-D) animals to yield heterozygous: heterozygous drd2-tdTomato (D2R) reporter mice. Sprague Dawley rats (P4-11; male and female) were injected i.p. with 10 μL (20 μg/μL dissolved in DMSO) of the retrograde tracer, Fast DiI (1,1′-dilinoleyl-3,3,3′,3′-tetramethylindocarbocyanine, 4-chlorobenzenesulphonate, D7756, Invitrogen) to back label parasympathetic preganglionic neurons (PPNs) in the lumbosacral spinal cord ^38^. After 3-7 days of transport, animals were euthanized for slice preparation and whole-cell electrophysiology.

### Solutions and slice preparation

D2R reporter mouse and neonatal (P7-14) rat spinal cord slices were prepared in the same way. Animals were anaesthetized i.p. with ketamine hydrochloride (100 mg/kg) followed by decapitation and exsanguination. A dorsal laminectomy was performed on ice to remove the lumbosacral spinal cord (L6-S1). Dura mater and dorsal/ ventral roots were trimmed in a Sylgard dish before the spinal cord was glued to a 4% agar block, ventral side down, and then to the specimen disc, with cyanoacrylate. Coronal sections (300 μm) were cut on a Vibratome VT1200s (Leica, Wetzlar) in ice-cold (2-4 °C) sucrose-substituted artificial cerebrospinal fluid pre-oxygenated with carbogen (sACSF; 350 mOsm/kg), containing (in mM); 250 Sucrose, 25 NaHCO_3_, 10 D-Glucose, 2.5 KCl, 1 NaH_2_PO_4_, 6 MgCl_2_, 1 CaCl_2_. Slices were then transferred into a carbogenated NMDG-HEPES recovery solution (NMDG-aCSF; 300 mOsm/kg) for 2-15 min at 32 °C, containing (in mM); 93 NMDG, 30 NaHCO_3_, 20 HEPES, 25 D-Glucose, 2.5 KCl, 1.2 NaH2PO_4_, 2 Thiourea, 3 Na Pyruvate, 10 MgSO_4_, 1 CaCl_2_, 5 Na Ascorbate (Ting et al., 2018). Slices were then placed in Recording aCSF (RaCSF; 300 mOsm/kg) and left at RT before transferring slices into the recording bath; RaCSF containing (in mM): 125 NaCl, 25 NaHCO_3_, 10 D-Glucose, 3 KCl, 1.2 KH_2_PO_4_, 1.2 MgSO_4_ and 2 CaCl_2_. Fire-polished thick-walled Borosilicate glass pipettes (3-6 MΩ, BF150-86-10, Sutter Instruments, Novato, CA) were pulled using a P-1000 Flaming/Brown (Sutter instruments, Novato, CA) and filled with an internal solution; containing (in mM); 135 K-gluconate, 6 NaCl, 4 NaOH, 10 HEPES, 11 EGTA, 2 Mg-ATP, 0.3 Na_2_-GTP, 1 MgCl_2_, 1 CaCl_2_, pH 7.3, adjusted with KOH (293 mOsm/kg).

### Whole-cell electrophysiology

All recordings were made in open, whole-cell configuration under voltage-or current-clamp conditions. Signals were acquired using Multiclamp 700B amplifier and Axon Digidata 1550 (Axon Instruments, San Jose, CA). All recordings occurred in carbogenated RaCSF at 32 °C. Drugs were perfused (3-5 mL/min) over the slice in RaCSF. Dopamine hydrochloride (10-100 µM, H8502, Sigma), Capromorelin (10 nM, CP-424,391, Pfizer), GHSR receptor antagonists agonists, JMV2959 (1 µM, HY-U00433A, Medchem Express), YIL781 (100 nM, 3959, Tocris Bioscience), PLCβ inhibitor (10 μM, U73122, AB120998, Abcam) and thapsigargin (1 μM, TG, T9033, Sigma) were washed onto slices (3-5 mL/ min) and holding current (pA) was assessed relative to baseline with VHolding at −60 mV. Recordings in adult mouse spinal cord slices were conducted in the presence of tetrodotoxin (1 μM, TTX, 14964, Cayman Chemical). Neurons with an access resistance > 25 ΜΩ were excluded, as were neurons with more than 50 pA leak current. Under voltage clamp conditions access resistance change was monitored during the recording and neurons were excluded if access resistance changed > 20%. Electrophysiology data were collected and analyzed with pClamp 10 and Clampfit software, respectively (Molecular devices, San Jose, CA).

Analysis of changes in holding currents were compared between baseline and peak drug response regions (mean pA) using Clampfit software (Molecular devices, San Jose, CA). Paired student’s t-test was performed on holding currents (pA) at baseline and conferred drug peak response region. P values and statistical significance has been stated in figure legends. Data are reported as mean ±SD.

### Calcium imaging

D2R-tdTomato (D2R) reporter mice were anaesthetized with isoflurane, injected i.p. with 0.01 mL ketamine hydrochloride (100 mg/kg) and decapitated. Spinal cord was removed promptly by dorsal laminectomy. 300 μm thick slices of lumbosacral spinal cord were cut in an ice-cold sucrose-aCSF. Spinal cord slices were left to recover in an NMDG-Hepes solution at 32 ℃ for 5-10 min followed by incubation in RaCSF at RT for 1 h. Spinal cord slices were incubated with Calbryte 520 AM (5 μM in RaCSF, 20650, AAT Bioquest) for 30-300 min at RT. For imaging, D2R-tdTomato labeled cells in the IML (intermediolateral nucleus) were located and time-lapsed imaged at 1.5 Hz using a 488-laser with a Zeiss LSM900 Confocal microscope at x20 objective (0.5NA). Baseline was recorded (2-5 min), followed by dopamine (30 μM in RaCSF) wash on (8-10 mins) and drug wash off (3-5 min) at a flow rate of 2-3 ml/min at 32 °C. To confirm if cells were loaded with Calbryte 520 AM, a high [K^+^] solution was washed onto the slices at the end of the recording (5-7 min; 76.8 NaCl, 75 KCl, 1.2 MgCl_2_, 2 CaCl_2_, 1.2 NaH_2_PO_4_, 14.4 NaHCO_3_, 11 D-Glucose). In calcium imaging experiments processing of files was carried out using FIJI (Schindelin et al., 2012).

Cytoplasmic regions of interest were highlighted, and the mean pixel intensity at each frame was measured and plotted as fluorescence versus time (z-profile). Data was converted to a relative scale (ΔF/ F0 baseline) and analyzed in Clampfit 10.4 (Molecular Devices, CA). Data is reported as mean ±SD. Calcium-dye-loaded D2R-tdTomato neurons were classified based on their relative fluorescence increase to K+ solution relative to the drug wash off period. An arbitrary threshold value of + 0.2 (ΔF/ F0) was used across spinal cord slices and experimental animals to exclude non-responders. All data were analyzed using GraphPad Prism (GraphPad, CA).

### Constructs

Our initial construct design was produced by Twist Bioscience in their ENTR vector, briefly, a synthetic construct encoding a CD-33 signal peptide followed by a FLAG epitope tag, SGGGGS linker and human GHSR was followed by an encephalomyocarditis virus internal ribosome entry site, which was then followed by a HA signal peptide, double cMyc epitope tag, SGGGGS linker and human DRD2 (long isoform, which is the post-synaptic form ^39^). Further constructs were generated in pENTR plasmid vectors by restriction digests and DRD2 only constructs were preceded by the internal ribosome entry site to achieve similar translational efficiency as the dual construct. Subsequently, LR recombination reactions were performed to shuttle entry vectors into pEF5/FRT/V5-DEST Gateway destination vector (Invitrogen) as previously described ^40^. The final constructs designed were spFLAG-GHSR1a-IRES-spcMyc-DRD2, spFLAG-GHSR1a, and IRES-spcMyc-DRD2. Monarch® DNA Gel Extraction Kits (NEB) were used per manufacturer’s instructions. DNA was produced by transforming DH5α competent cells by selection with 50 µg/mL kanamycin for entry vectors and carbenicillin for destination vectors. Maxipreps (Qiagen) and minipreps (Promega) were used to purify DNA as per manufacturer’s instructions. Generated vectors were verified by Sanger sequencing in forward and reverse directions.

### Cell line generation and cell culture

FlpIn CHO and HEK293 cell lines were purchased from Invitrogen and maintained in Dulbecco’s Modified Eagle’s Medium supplemented with 5% Foetal Bovine Serum (ThermoFisher Scientific (Gibco)) at 37°C. Expression constructs (pEF5/FRT/V5-DEST) and pOG-44 Flp Recombinase Expression Vector (ThermoFisher Scientific (Invitrogen)) were transfected into target cells using Lipofectamine2000 (Invitrogen) per manufacturer’s instructions and stable cell lines were generated by polyclonal selection in 600 µg/ml hygromycin (InvivoGen).

### Fluorescently labeled ligands

Full-length human ghrelin labeled with Alexa Fluor 568 (ghrelin-AF568) was synthesized and validated as described previously (Liu et al., 2020). The DRD2 specific fluorescent ligand (ZX-1) was generated by synthetic chemistry based on the known binding pose of haloperidol in DRD2 following previously published methods ^41^ for generating haloperidol analogues and incorporating a linker predicted to face the extracellular space to allow attachment of sulfo-Cy5 (synthesized in-house and described below, synthesis in Supp Fig. 4D).

#### General Chemistry

All solvents and chemicals were purchased from standard suppliers and were used without any further purification. ^1^H NMR and ^13^C NMR spectra were acquired at 400.13 and 100.62 MHz, respectively, on a Bruker Avance III Nanobay 400 MHz NMR spectrometer coupled to the BACS 60 automatic sample changer and equipped with a 5 mm PABBO BB-1H/ D Z-GRD probe. All spectra obtained was processed using MestReNova software (v.6.0). Chemical shifts (δ) for all ^1^H NMR spectra are reported in parts per million (ppm) using tetramethylsilane (TMS, 0 ppm) as the reference. The data for all spectra are reported in the following format: chemical shift (δ), (multiplicity, coupling constants *J* (Hz), integral), where the multiplicity is defined as s = singlet, d = doublet, t = triplet, q = quartet, p = pentet, and m = multiplet. ^13^C NMR were routinely carried out as *J*-modulated spin-echo experiments (JMOD), all ^13^C NMR δ are reported in ppm. Thin layer chromatography (TLC) was carried out routinely on silica gel 60F_254_ precoated plates (0.25 mm, Merck). Flash column chromatography was carried out using Davisil LC60A silica gel, 40-63 μm. The PREP HPLC is an Agilent 1260 infinity coupled with binary prep pump and Agilent 1260 FC-PS fraction collector. The column is an Alltima C8 5 μm, 22 mm × 250 mm. The preparative HPLC operates on Agilent OpenLAB CDS Rev C.01.04 software. Solvent A is water + 0.1% TFA (or 0.1% CH_3_COOH if compounds are unstable under TFA) and solvent B is acetonitrile + 0.1% TFA (or 0.1% CH_3_COOH if compounds are unstable under TFA). Samples were run using various gradient methods (5-100% solvent B over 10-30 min).

Liquid chromatography mass spectrometry (LCMS) was performed on one of two instruments; either an Agilent 6100 Series Single Quad LC/MS or an Agilent 1200 Series HPLC (equipped with a 1200 Series G13111A Quaternary Pump, G1329A Thermostatted Autosampler, and a G1314B Variable Wavelength Detector) and the data was processed using LC/MSD Chemstation Rev.B.04.01 SP1 coupled with Easy Access Software. Both systems were equipped with a Reverse Phase Luna C8(2) (5 μm, 50 × 4.6 mm, 100 Å) column maintained at 30°C. An MeCN gradient (5-100%) was used to obtain optimal separation, where 4 min were required for the gradient to reach 100% MeCN and maintained for a further 3 minutes before requiring 3 min to return to the initial gradient of 5% MeCN (total run time = 10 min). Solvent A = 0.1% aqueous formic acid; Solvent B = MeCN/ 0.1% formic acid.

The purity and retention time of final products were determined using analytical HPLC and high-resolution mass spectrometry (HRMS). Analytical HPLC was carried out using an Agilent 1260 Infinity Analytical HPLC fitted with a Zorbax Eclipse Plus C18 Rapid Resolution column (100 mm × 4.60 mm, 3.5 μm) using a binary solvent system: solvent A of 0.1% aqueous TFA; solvent B of 0.1% TFA in MeCN. Gradient elution was achieved over 10 min using 95% A + 5% B to 100% B over 9 min, and 100% B maintained for 1 minute at a flow rate of 1 mL/min monitored at both 214 and 254 nm. HRMS were conducted on an Agilent 6224 TOP LC/MS Mass Spectrometer coupled to an Agilent 1290 Infinity. All data was acquired and reference mas corrected *via* dual-spray electrospray ionisation (ESI) source. Each scan or data point on the total ion chromatogram (TIC) is average of 13700 transients, producing one spectrum per second. Mass spectra were created by averaging the scans across each peak and background subtracted against the first 10 seconds of the TIC. Data acquisition was carried out using the Agilent Mass Hunter Data Acquisition software version B.05.00 Build 5.0.5042 and analysis was performed using Mass Hunter Qualitative Analysis version B.05.00 Build 5.0.519.13.

### 4-(4-(4-Bromophenyl)-4-hydroxypiperidin-1-yl)-1-(4-fluorophenyl)butan-1-one (4)

To a 5 mL vial equipped with a stirrer bar was added 4-chloro-4′-fluorobutyrophenone (**2**, 500 mg, 410 μL, 2.5 mmol, 1.0 equiv.), 4-(4-bromophenyl)-4-hydroxypiperidine (**3**, 1.28 g, 5.0 mmol, 2.0 equiv.) and anhydrous KI (12.4 mg, 74.8 μmol, 0.03 equiv.). The vial was sealed after 2 mL of anhydrous toluene was added. The mixture was stirred at 130°C for 24 h. After cooling to room temperature, the reaction mixture was diluted with sat. NaHCO_3_ (20 mL) and the aqueous layer was extracted with EtOAc (3 × 20 mL). The combined organic layers were washed with brine, dried over anhydrous MgSO_4_, filtered, and concentrated. The residue was purified by flash chromatography (DCM/MeOH/Et_3_N, 20:1:0.2) and provided the title compound (570 mg, 54% yield) as a white solid. ^1^Η ΝΜR (DMSO-*d*_6_) *δ* 8.10-8.05 (m, 2H), 7.48-7.45 (m, 2H), 7.39-7.33 (m, 2H), 7.31-7.28 (m, 2H), 4.82 (s, 1H), 2.97 (t, *J* = 6.8 Hz, 2H), 2.57 (d, *J* = 10.8 Hz, 2H), 2.35 (t, *J* = 6.7 Hz, 2H), 2.31 (t, *J* = 9.2 Hz, 2H), 1.83 (p, *J* = 6.8 Hz, 2H), 1.65 (td, *J* = 12.7 Hz, 4.1 Hz, 2H), 1.45 (d, *J* = 12.1 Hz, 2H). ^13^C NMR DEPTQ (DMSO-*d*_6_) *δ* 198.3, 166.1, 163.6, 149.6, 134.0, 130.9, 130.8, 130.6, 127.2, 119.2, 115.7, 115.5, 69.6, 57.3, 48.9, 37.7, 35.7, 22.0. LCMS (ESI): *m/z* 419.8 and 421.8. Analytical HPLC: >95%, *t*_R_ = 5.351 min.

### *tert*-Butyl (3-(4-(1-(4-(4-fluorophenyl)-4-oxobutyl)-4-hydroxypiperidin-4-yl)phenyl)prop-2-yn-1-yl)carbamate (6)

Dry DMF and Et_2_NH were degassed (30 min) with N_2_. **4** (500 mg, 1.19 mmol, 1 equiv.), Pd(PPh_3_)_2_Cl_2_ (41.8 mg, 59.5 μmol, 0.05 equiv.), CuI (11.3 mg, 59.5 μmol, 0.05 equiv.), triphenylphosphine (62.4 mg, 238 μmol, 0.2 equiv.), *N*-*Boc*-propargylamine (**5**) (554 mg, 3.57 mmol, 3 equiv.), dried diethylamine (3.5 mL) and DMF (1.5 mL) were added into a microwave reaction vial quickly, degassed with N_2_ again, and then stirred under N_2_ at 120°C for 40 min in the microwave cavity. The reaction mixture was diluted with saturated NaHCO_3_ solution (20 mL) and extracted with EtOAc (3 × 20 mL). Then the combined organic phase was washed with brine, dried over anhydrous MgSO_4_ and then concentrated under reduced pressure. The crude mixture was purified by flash column chromatography (DCM/methanol/Et_3_N, 20:1:0.1). The relevant fractions were combined and reduced *in vacuo* to afford a light brown solid (377 mg, 64%). ^1^Η ΝΜR (CDCl_3_) *δ* 7.99 (dd, *J* = 8.7 Hz, 5.5 Hz, 2H), 7.38-7.33 (m, 4H), 7.11 (t, *J* = 8.6 Hz, 2H), 4.94 (s, 1H), 4.10 (s, 2H), 2.96 (t, *J* = 7.0 Hz, 2H), 2.78 (d, *J* = 10.9 Hz, 2H), 2.46 (dd, *J* = 9.3 Hz, 16.8 Hz, 4H), 2.04-1.93 (m, 4H), 1.66 (d, *J* = 12.7 Hz, 2H), 1.44 (s, 9H). ^13^C NMR DEPTQ (CDCl_3_) *δ* 198.5, 167.0, 155.5, 148.8, 133.7, 133.7, 131.7, 130.8, 130.7, 124.7, 121.3, 115.8, 115.6, 85.4, 83.0, 77.5, 77.2, 76.8, 71.2, 57.9, 50.7, 49.4, 38.2, 36.3, 28.6, 28.5, 21.8. LCMS (ESI): *m/z* 495.1. Analytical HPLC: >95%, *t*_R_ = 5.634 min. HRMS (ESI) TOF (*m/z*): [M+H]+ 495.2654 calcd for C_29_H_35_FN_2_O_4_; found [M+H]^+^ 495.2654

### 4-(4-(4-(3-Aminoprop-1-yn-1-yl)phenyl)-4-hydroxypiperidin-1-yl)-1-(4-fluorophenyl)butan-1-one (7)

To a 5mL RBF containing **6** (117 mg, 0.240 mmol, 1 equiv.) was added 4.0 M HCl in dioxane solution (1.6 mL) at room temperature, and the reaction mixture was stirred under nitrogen atmosphere for 2.5 h. The reaction was quenched with saturated NaHCO_3_ solution and then extracted with DCM (3 x 5 mL). The combined organic layer was washed with brine, dried with anhydrous MgSO_4_ and then concentrated *in vacuo*. The crude mixture was purified flash column chromatography (DCM/MeOH/NH_3_, 4:1:0.1). The relevant fractions were combined and reduced *in vacuo* to afford the free amine as a white solid (25 mg, 21%). ^1^Η ΝΜR (CD_3_OD) *δ* 8.12-8.07 (m, 2H), 7.53 (q, *J* = 8.6 Hz, 4H), 7.27-7.22 (m, 2H), 4.04 (s, 2H), 3.61 (d, *J* = 12.1 Hz, 2H), 3.46 (td, *J* = 12.4 Hz, 1.9 Hz, 2H), 3.30-3.28 (m, 1H), 3.27 (d, *J* = 5.4 Hz, 1H), 3.23 (t, *J* = 6.7 Hz, 2H), 2.34 (td, *J* = 14.4 Hz, 4.2 Hz, 2H), 2.24-2.16 (m, 2H), 2.01 (s, 1H), 1.98 (s, 1H). ^13^C NMR DEPTQ (CD_3_OD) *δ* 198.7, 168.6, 166.1, 149.6, 134.5, 133.0, 132.1, 132.0, 126.1, 121.9, 116.8, 116.6, 87.4, 81.6, 69.4, 57.8, 50.3, 36.6, 35.9, 30.7, 19.6. Analytical HPLC: >95%, *t*_R_ = 4.041 min. HRMS (ESI) TOF (*m/z*): [M+H]^+^ 395.2129 calcd for C_24_H_27_FN_2_O_2_; found [M+H]^+^ 395.2134.

### *tert*-Butyl(6-((3-(4-(1-(4-(4-fluorophenyl)-4-oxobutyl)-4-hydroxypiperidin-4-yl)phenyl)prop-2-yn-1-yl)amino)-6-oxohexyl)carbamate (9)

Compound **8** (88.0 mg, 0.380 mmol, 1.5 equiv.) was dissolved in DMF (2 mL). PyBOP (198 mg, 0.380 mmol, 1.5 equiv.) and DBU (116 mg, 0.760 mmol, 113 μL, 3 equiv.) were added into the vial. After 5 minutes, **7** (100 mg, 0.253 mmol, 1 equiv.) was dissolved in DMF in another vial and transferred into the reaction mixture. The reaction was stirred at room temperature for 8 h. The reaction mixture was concentrated under reduced pressure and purified by flash column chromatography (DCM/MeOH/NH_3_, 20:1:0.1). The relevant fractions were combined and concentrated *in vacuo* to afford light brown solid (110 mg, 71%). ^1^Η ΝΜR (CD_3_OD) *δ* 8.12-8.07 (m, 2H), 7.44 (dd, *J* = 6.8 Hz, 2.0 Hz, 2H), 7.39 (dd, *J* = 6.8 Hz, 2.0 Hz, 2H), 7.24 (tt, *J* = 8.8 Hz, 2.0 Hz, 2H), 4.18 (s, 2H), 3.16 (dd, *J* = 14.4 Hz, 7.2Hz, 2H), 3.12 (t, *J* = 6.8 Hz, 2H), 3.02 (t, *J* = 7.0 Hz, 2H), 2.87 (t, *J* = 11.4 Hz, 2H), 2.79 (t, *J* = 7.2 Hz, 2H), 2.23 (t, *J* = 7.5 Hz, 2H), 2.16-2.03 (m, 4H), 1.79 (d, *J* = 12.9 Hz, 2H), 1.64 (p, *J* = 7.6 Hz, 2H), 1.49-1.47 (m, 2H), 1.42 (s, 9H), 1.39-1.33 (m, 2H). ^13^C NMR DEPTQ (DMSO-*d*_6_) *δ* 197.9, 171.9, 166.1, 163.7, 155.6, 149.8, 143.0, 133.7, 130.9, 130.8, 125.0, 120.3, 115.7, 115.5, 86.8, 81.4, 77.3, 69.1, 56.5, 39.7, 36.6, 35.5, 35.1, 29.3, 28.5, 28.3, 28.3, 26.0, 24.9, 20.7. LCMS (ESI): *m/z* 608.1. Analytical HPLC: 94%, *t*_R_ = 5.527 min. HRMS (ESI) TOF (*m/z*): [M+H]^+^ 608.3494 calcd for C_35_H_46_FN_3_O_5_; found [M+H]^+^ 608.3511.

### 6-Amino-N-(3-(4-(1-(4-(4-fluorophenyl)-4-oxobutyl)-4-hydroxypiperidin-4-yl)phenyl)prop-2-yn-1-yl)hexanamide (10)

To the protected amine **9** (48 mg, 79.0 µmol, 1 equiv.) was added an excess of 4.0 M HCl/1,4-dioxane solution (1 mL). The reaction mixture was stirred at room temperature overnight. The solvent was removed under reduced pressure. The crude mixture was purified flash column chromatography (DCM/MeOH/NH_3_, 4:1:0.1). The relevant fractions were combined and reduced *in vacuo* the free amine as a white solid (22 mg, 55%). ^1^Η ΝΜR (DMSO-*d*_6_) *δ* 8.39 (t, *J* = 5.4 Hz, 1H), 8.09-8.05 (m, 2H), 7.36 (t, *J* = 8.8 Hz, 2H), 7.32 (t, *J* = 4.8 Hz, 4H), 4.10 (d, *J* = 5.3 Hz, 2H), 2.96 (t, *J* = 6.7 Hz, 2H), 2.65 (t, *J* = 7.3 Hz, 2H), 2.56 (d, *J* = 10.1 Hz, 2H), 2.34 (t, *J* = 6.4 Hz, 2H), 2.31 (t, *J* = 10.8 Hz, 2H), 2.10 (t, *J* = 7.4 Hz, 2H), 1.83 (p, *J* = 6.8 Hz, 2H), 1.64 (td, *J* = 12.8, 3.8 Hz, 2H), 1.56-1.51 (m, 2H), 1.48 (d, *J* = 7.7 Hz, 2H), 1.46-1.43 (m, 2H), 1.26 (p, *J* = 8.0 Hz, 2H). ^13^C NMR DEPTQ (DMSO-*d*_6_) *δ* 198.3, 171.8, 166.1, 163.6, 150.7, 134.0, 130.9, 130.8, 125.1, 120.0, 115.7, 115.5, 86.7, 81.5, 69.7, 57.3, 48.9, 37.6, 35.6, 35.0, 28.7, 28.5, 25.7, 24.8, 22.1. Analytical HPLC: >95%, *t*_R_ = 4.239 min. HRMS (ESI) TOF (*m/z*): [M+H]^+^ 508.2970 calcd for C_30_H_38_FN_3_O_3_; found [M+H]^+^ 508.2992.

### 1-(6-((6-((3-(4-(1-(4-(4-Fluorophenyl)-4-oxobutyl)-4-hydroxypiperidin-1-ium-4-yl)phenyl)prop-2-yn-1-yl)amino)-6-oxohexyl)amino)-6-oxohexyl)-3,3-dimethyl-2-((1E,3E)-5-((E)-1,3,3-trimethyl-5-sulfoindolin-2-ylidene)penta-1,3-dien-1-yl)-3H-indol-1-ium-5-sulfonate 2,2,2-trifluoroacetate (1)

Sulfo-cyanine5 carboxylic acid **11** (3.45 mg, 6.80 µmol, 1 equiv.) was dissolved in DMF (0.6 mL) in an Eppendorf vial. PyBOP (4.43 mg, 8.50 µmol, 1.5 equiv.) and DIPEA (2.20 mg, 17.0 µmol, 3 equiv.) were added quickly into the vial. Following this, **10** (3.45 mg, 6.80 µmol, 1.2 equiv.) was dissolved in DMF (0.3 mL) in another vial and transferred into the sulfo-cyanine5 carboxylic acid solution. The reaction was stirred at room temperature for 12 h in the absence of light. The reaction mixture was monitored by LC-MS and then purified immediately *via* preparative-HPLC. The clean fractions were collected, pooled and the water removed *via* lyophilization to obtain the product as the TFA salt (0.66 mg, 7.8%). ^1^Η ΝΜR (DMSO-*d*_6_) *δ* 9.17 (s, 1H), 8.38-8.31 (m, 3H), 8.11-8.07 (m, 2H), 7.82 (dd, *J* = 2.6 Hz, 1.6 Hz, 2H), 7.73 (t, *J* = 6.2 Hz, 1H), 7.66 (dd, *J* = 4.7 Hz, 1.5 Hz, 1H), 7.64 (dd, *J* = 4.7 Hz, 1.6 Hz, 1H), 7.39 (t, *J* = 8.9 Hz, 2H), 7.34-7.27 (m, 2H), 6.56 (t, *J* = 12.4 Hz, 1H), 6.30 (d, *J* = 14.2 Hz, 1H), 6.25 (d, *J* = 14.1 Hz, 1H), 5.54 (s, 1H), 4.09 (d, *J* = 5.3 Hz, 4H), 3.60 (s, 3H), 2.99 (dd, *J* = 12.5 Hz, 6.6 Hz, 2H), 2.67 (q, *J* = 2.0 Hz, 2H), 2.56-2.54 (m, 2H), 2.54-2.52 (m, 2H), 2.48-2.47 (m, 2H), 2.46-2.45 (m, 2H), 2.33 (q, *J* = 1.6 Hz, 2H), 2.12-2.07 (m, 4H), 2.03 (t, *J* = 7.6 Hz, 2H), 1.78 (d, *J* = 13.1 Hz, 2H), 1.70-1.67 (m, 2H), 1.65 (s, 6H), 1.63 (s, 6H), 1.56-1.54 (m, 4H), 1.37-1.32 (m, 2H), 1.24-1.21 (m, 2H). Analytical HPLC: >95%, *t*_R_ = 5.167 min. HRMS (ESI) TOF (*m/z*): [M+H]^+^ 1132.4934 calcd for C_62_H_74_FN_5_O_10_S_2_; found [M+H]^+^ 1132.4897.

### Saturation ligand binding

To determine ligand affinity, no wash binding assay was performed. FlpIn CHO cells expressing different GPCRs were detached from cell culture flasks using PBS (Phosphate Buffered Saline, pH 7.4) containing 5.3 mM EDTA and then chilled on ice before being transferred to wells of a 96-well tray in 100 µL at ∼6×10^4^ cells per mL. All subsequent centrifuge steps were carried out at 4°C for 3 minutes at 400 ×g. Cells were washed twice with HBSS binding buffer (Hank’s Balanced Salt Solution containing 0.05% Ovalbumin, 20 mM HEPES, and 0.05% Sodium Azide), by centrifugation followed by supernatant aspiration, gentle resuspension, and blocking in the same buffer for additional 20 minutes. ZX-1 dilutions ranging from 0.316 nM to 100 nM and ghrelin:AF568 dilutions ranging from 0.1 nM to 100 nM were prepared in binding buffer.

Following blocking step, cells were centrifuged, supernatant removed and resuspended in 200 µL of the appropriate ligand concentration and an incubated in the dark at 4°C for 4 h. Cells were centrifuged and allowed to remain in binding buffer with ligand. Immediately prior to loading into the flow cytometer the supernatant was carefully aspirated and cells resuspended in 200 µL of binding buffer containing 500 nM Sytox Blue (ThermoFisher Scientific (Invitrogen)) and analyzed by BD Fortessa cytometer (BD Biosciences). Raw data was processed initially on FlowJo (Becton, Dickinson and Company, version 10.6.2) to gate single, live cells. Mode values for each sample was then converted to a fold shift over baseline (unstained cell sample) and entered into PRISM 9 (GraphPad). Affinity and Bmax values were determined for each individual experiment using a one-site (Total and nonspecific binding) model in PRISM 9 (GraphPad):

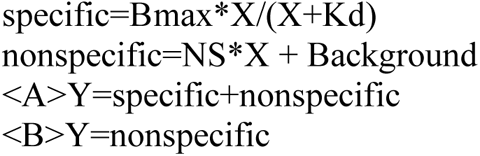

Individual values were used to perform student’s T-test and statistical significance was deemed where p <0.05 and is denoted by asterisks throughout. Pooled data was plotted as mean ±SD binding curves for visualization purposes.

### Flow cytometry for cell surface expression

M1 anti-FLAG antibody was produced in-house from 4E11 hybridoma (ATCC: HB-9259) by purification from the hybridoma supernatant using custom FLAG peptide column (DYKDDDDK-GGSC (custom GL Biochem) coupled to SulfoLink resin (ThermoFisher Scientific, cat# 20402)) and confirmed by gel electrophoresis. Anti-cMyc antibody was produced in-house from 9E10 hybridoma (ATCC: CRL-1729) by purification from the hybridoma supernatant using Protein-G Sepharose column (Cytiva, cat# 17040501) and confirmed by gel electrophoresis. M1 anti-FLAG (4E11) and anti-cMyc (9E10) antibodies were conjugated to AF568 and AF647, respectively, as described previously ^42^.

One-layer staining: cells were harvested using PBS containing 5.3 mM EDTA, chilled on ice, and 100 µL per well of ∼6×10^4^ cells per mL transferred to a v-bottom 96-well tray. All centrifuge steps were carried out at 4°C, 3 minutes at 400 ×g. Cells were washed twice with HBSS buffer (containing 0.05% ovalbumin, 20 mM HEPES, and 0.05% Sodium Azide) by centrifugation followed by aspirating supernatant and gentle resuspension in buffer. Washes were followed by blocking in the same buffer for 20 minutes on ice. Following blocking, cells were stained in 200 µl HBSS buffer containing 1 µg/ml 4E11:AF568 and 9E10:AF647 for two hours. Cells were washed once with HBSS buffer then centrifuged again before being resuspended in 200 µL HBSS buffer containing 500 nM Sytox Blue (ThermoFisher Scientific (Invitrogen)) before being analyzed by BD Fortessa cytometer (BD Biosciences).

Two-layer staining: as above with unconjugated primary antibodies. After primary antibody stain (separately for each antibody) and 2× washes, cells were stained with 2 µg/ml AF647-conjugated secondary antibody (Thermo Fisher Scientific, cat# A-21236) for one h. Cells were then washed once with 1 ml of HBSS buffer before staining with. Cells were washed once with HBSS buffer then centrifuged again before being resuspended in 200 µL HBSS buffer containing 500 nM Sytox Blue (ThermoFisher Scientific (Invitrogen)) before being analyzed on a Stratadigm cytometer.

Raw data was processed initially on FlowJo (Becton, Dickinson and Company, version 10.6.2) to gate single, live cells and provide representative histograms. Mode values for each sample was then converted to a fold shift over baseline (unstained cell sample) and entered into PRISM 9 (GraphPad) to allow statistical comparison between cell lines.

### Calcium assay

Cells were harvested and seeded to a black 96-well plate (Corning, cat# 3916) at a density of 3×10^4^ cells/well and incubated overnight (1×10^4^ 1 for 384 well plates (Corning, cat# CLS3764)). The next day, media was aspirated, and cells were washed with 100 µL calcium buffer (1×x HBSS buffer, 20 mM HEPES (pH 7.4), 4 mM Probenecid, 0.1% ascorbic acid, and 0.05% (w/v) OVA).

Cells were then incubated at 37°C for 45 min in 90 µl calcium buffer (40 µl for 384-well plates) containing 1 µM Fluo-8 AM (Abcam, cat# 142773). 10x (5x for 384-well plates) drug dilutions were prepared in calcium assay buffer. YIL781 (Tocris, cat# 3959), sulpiride (Tocris, cat# 0894), and YM254890 (Tocris, cat# 7352) were added to cells 30 min prior to reading. Pertussis toxin (Tocris, cat# 3097) was added 16 h prior. Plates were read (excitation at 480 nm and emission at 542 nm) on the PHERAstar (BMG Labtech, 96-well), µFDSS (Hamamtsu, 96-well) or FLIPR Tetra (Molecular Devices, 384-well). For 96-well plates readings were taken every 3 seconds in the PHERAstar, every 1 s in the µFDSS and for 384-well plates they were taken every 0.5 seconds on a FLIPR. For analysis, vehicle and baseline responses were subtracted and maximum responses were plotted on PRISM 9 software (GraphPad), normalized to maximal ionomycin response, and fitted to a log (agonist) vs response (three parameter) curve.

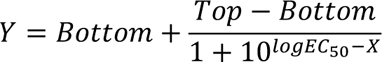

### cAMP inhibition assay

Cells were harvested and seeded to a black 96-well plate (Corning, cat# 3917) at a density of 3×10^4^ cells/well in 90 μL of their regular media and incubated overnight. Approximately 24 hours prior to assay, transfection mix (30 ng CAMYEN biosensor/well) (Jiang et al., 2007) was prepared in Opti-Mem (Gibco) with DNA to PEI MAX (Polysciences, cat# 24765) ratio of 1:6 (µg:µg) per the manufacturer’s instructions and incubated at room temperature for 30 minutes. Following incubation, 10 µl of transfection mix was added to each well and incubated overnight. The next day, media was aspirated, and cells were washed with 100 µL cAMP assay buffer (1x HBSS buffer, 20 mM HEPES (pH 7.4), 0.1% ascorbic acid, and 0.05% (w/v) OVA). Cells were then incubated at 37°C for 45 minutes in 80 µL cAMP assay buffer. All drug dilution series, inhibitors, and forskolin were made in cAMP buffer at 10x concentrations. YIL781 (Tocris, cat# 3959), sulpiride (Tocris, cat# 0894), and YM254890 (Tocris, cat# 7352) were added to cells 30 min prior to reading.

Pertussis toxin (Tocris, cat# 3097) was added 2 hours following transfection (approximately 20 hours prior to assay). Five min prior to reading, 10 μL of Furimazine (TargetMol, cat# T15359, final concentration of 1 μM) was added to each well. Plates were read on the PHERAstar (BMG Labtech) plate reader at 37°C and CAMYEN dual emission was detected at 475 nm and 525 nm for 22.5 min (90 cycles, 15 s/cycle). Reading included 10 cycles for baseline recording followed by cAMP activation by forskolin addition (3 µM) for 30 cycles and cAMP inhibition by stimulation with DRD2 agonists for further 30 cycles. For analysis, vehicle and baseline responses were first subtracted to obtain the corrected forskolin activation curves. The final 6 last points of each curve were then used to obtain the baseline correction of cAMP inhibition followed by vehicle subtraction. The Area Under the Curve (AUC) was calculated using PRISM 9 software (GraphPad), plotted, and fitted to a log (inhibitor) vs response (three parameter) curve.

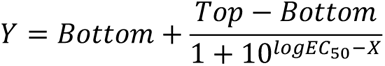

### TRUPATH Assay

Cells were seeded into white 96-well plates (Corning, cat# 3917) at a density of 22,000 cells per well. 12 h after seeding, cells were transfected using a DNA ratio of 1:1:1 for Gα-RLuc8, Gβ, and GFP2-Gγ constructs (100 ng total DNA per well). PEI MAX (Polysciences, cat# 24765) was used for transfection at a ratio of 1:4 (DNA: PEI, µg:µg). DNA and PEI were diluted with an equal volume of Opti-Mem (Gibco) separate tubes then the PEI mix was added to the DNA (vortexed gently), resulting in a 10x transfection mix. Following a 30-min incubation at room temperature, the transfection mixture was then diluted in media to a final concentration of 1 ng/µL DNA. Media was aspirated from cells and replaced with 100 μL per well of the transfection mix and incubated (37°C, 5% CO_2_) for 24 h. On the day of the experiment, a fresh buffer (20 mM HEPES buffer, 0.05% ovalbumin, 1x HBSS buffer, and 0.1% ascorbic acid) with a pH of 7.4 was prepared. The media was aspirated and cells washed once with 100 μL of buffer. Cells were then incubated for 20 min at 37°C in 80 μL of buffer per well. After the 20-min incubation, 10 μL of 10x prolume purple in buffer (Nanolight #369, final concentration of 6.7 μM) was added to each well and incubated for 10 min in the PHERAstar (BMG Labtech) plate reader at 37°C. 10x drug stocks (Dopamine, Pramipexole, Ghrelin, and Capromorelin) were prepared using the same buffer and brought to 37°C. Measurement was done half of the plate at a time on the PHERAstar (BMG Labtech) plate reader. BRET2 plus dual emission was detected at 410/80 nm and 515/30 nm for 12 min (60 cycles, 12 s/cycle). Reading included 10 cycles of baseline followed by stimulation with 10 μL of the 10x drugs for further 50 cycles. For analysis the BRET ratio (515/410) was calculated and the vehicle and baseline responses were subtracted to obtain the corrected signal curve. AUC was calculated by PRISM 9 software (GraphPad), plotted, and fitted to a log (inhibitor) vs response (three parameter) curve.

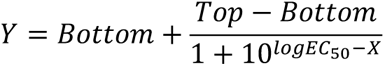

### Deconvolution Microscopy

Cells were harvested and plated at 2×10^4^ per well to an 8 well IBIDI μ-slide (Ibidi GmbH) with polymer coverslip and IBIDI coat and cell media to incubate at 37 °C overnight. Antibodies were diluted fresh the next day for staining. Primary antibody staining solutions were 10 μg/mL 4E11:AF568 and 3 μg/mL 9E10:AF647 in 1x HBSS, 20mM HEPES, pH=7.4 at 4°C (see above Flow Cytometry section for description of antibodies). Wash buffer: 1x HBSS, 20mM HEPES, 0.1% OVA, 0.02% sodium azide, pH=7.4, 4°C. Blocking buffer: 5% BSA, 1x HBSS, 20 mM HEPES, 0.1% OVA, 0.02% sodium azide, pH=7.4, 4°C. Cells were chilled 1 hr on ice, then washed 1x and blocked for 30 min on ice. Blocking buffer was aspirated and replaced with primary antibody and incubated for 90 min on ice. Cells were washed 4x 5 minutes and fixed for 1 minute on ice with ice cold 4% paraformaldehyde (PFA, from 8% methanol-free electron microscopy grade, Electron Microscopy Services) in 1x HBSS, 20mM HEPES. Fixing buffer was aspirated and cells washed 3x in wash buffer then stored overnight in fridge. The next day, cells were imaged on a Sp8 lightning confocal (Leica) with a 63x 1.30 NA, glycerol objective, lightning (deconvolution) mode for optimized resolution, 4x scan with averaging.

### STORM imaging

Cells were harvested and plated at a density of 1×10^5^ per well to an 8 well IBIDI μ-slide (Ibidi GmbH) with glass coverslip (coated with 5 μg/cm^2^ fibronectin (Roche) for an hour at room temperature) and cell media to incubate at 37°C overnight. Cells were chilled on ice and washed 3 times with cold microscopy buffer: 1x HBSS, 20mM HEPES, 0.05% OVA, 0.05% sodium azide, pH 7.4. The staining solution was prepared in microscopy buffer and contained 316 nM ZX-1 and 100 nM ghrelin:AF568. Following 3 hours of staining at 4°C, cells were washed 3 times with imaging buffer and fixed for 60 minutes at 4°C with buffer containing 1x HBSS, 20mM HEPES, 4% PFA (EMS), and 0.2% glutaraldehyde (EMS), pH 7.4. Following fixation, cells were washed 3 times and stored in the microscopy buffer at 4°C to be imaged within 4 hours. For STORM microscopy, buffer was aspirated and replaced with imaging buffer containing 50 mM Tris, 2.5 mM NaCl, 10% glucose, 10 mM MEA (mercapto ethylamine), 250 units/ml glucose oxidase (Sigma), 2500 units/ml catalase. Image acquisition was performed using a Nikon CFI Apo TIRF 100x/1.49 NA oil-immersion objective on a single-molecule localisation microscope (SAFe 360; Abbelight, France) built around a Nikon ECLIPSE Ti2 body and equipped with two ORCA-Fusion BT sCMOS cameras (Hamamatsu Photonics K.K., Japan) and controlled by Abbelight NEO software. Both fluorophores were bleached simultaneously with 561 and 640 laser channels (20 % laser power) for several minutes at 50 frames/second or until they started blinking. After bleaching, STORM images were acquired at 50 frames/second in HILO (highly inclined and laminated optical sheet) for 40000 frames while gradually increasing laser power for continuous blinking of the fluorophores.

### STORM analysis

Raw image stacks were analyzed with the ThunderStorm plugin in Fiji (ImageJ) using parameters matched to the SAFe 360 (97 nm pixel, 0.24 photoelectrons per A/D count, 0.95 base level A/D counts and a magnification of 6.466 to achieve an effective xy resolution of 15 nm). All transmitted stacks (far red) were analyzed first and drift corrected using a 9 iteration, model-based correction. The resulting drift table was then applied to the matched reflected channel, which was then processed. Resulting localization reconstructions were then analyzed for nearest neighbor distance using the following script:

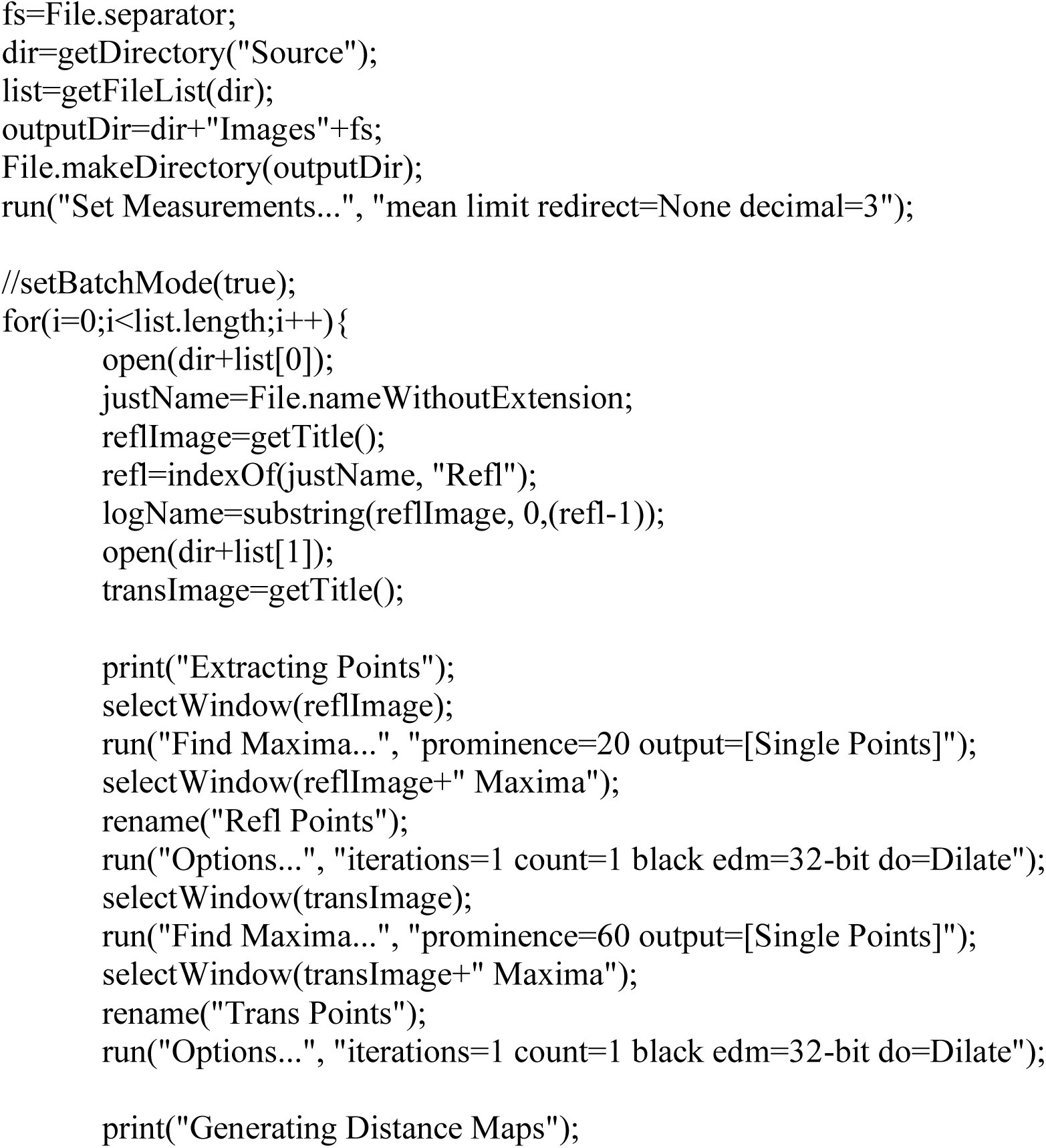
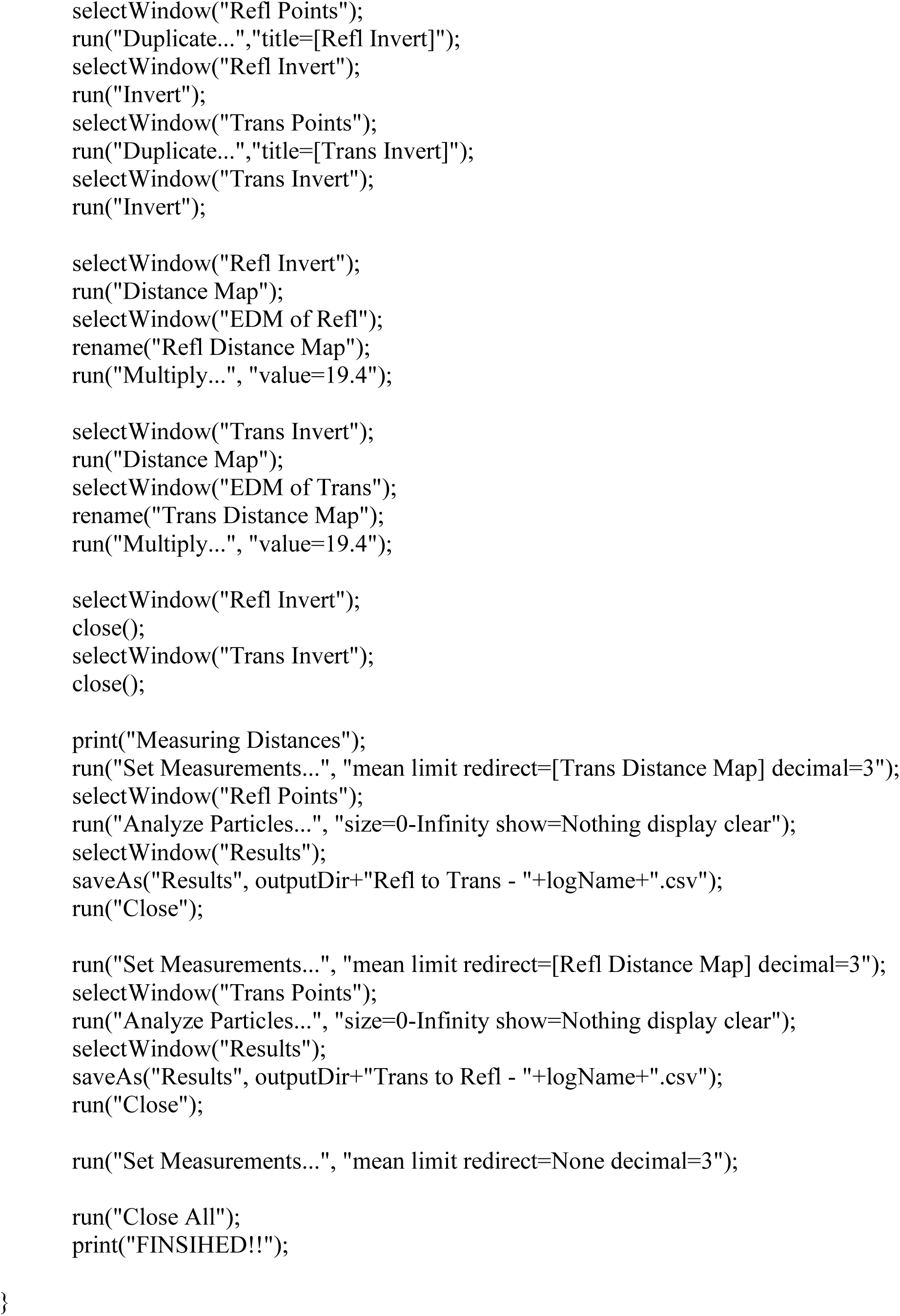

Nearest neighbor data was generated for each experiment; within experiment images were pooled with equal weighting into cumulative histograms with a bin width of 20 nm.

---

**Fig. S2.**
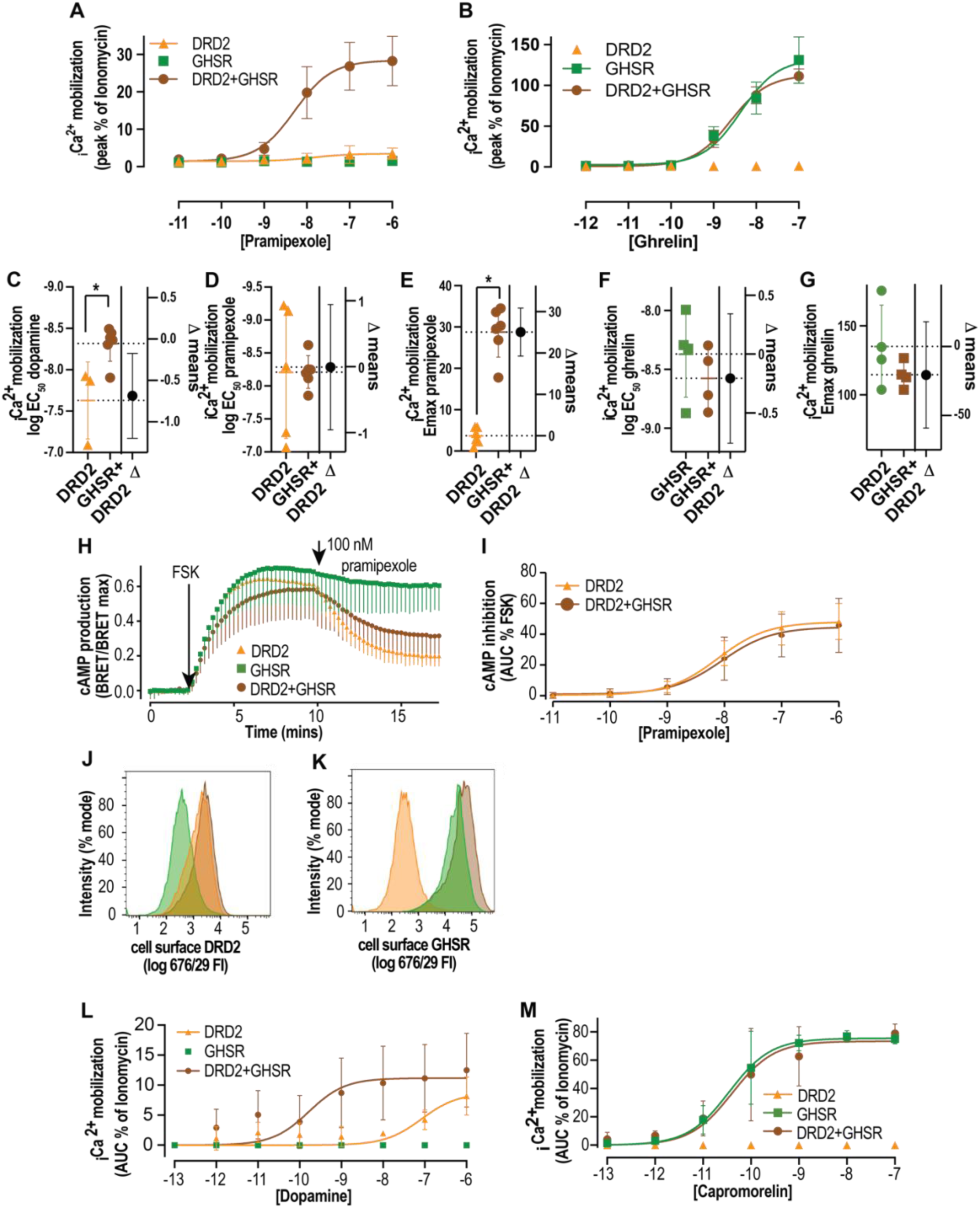
GHSR provides a pharmacological switch for DRD2 in recombinant cell lines. (**A**) GHSR expression is permissive for _i_[Ca^2+^] of DRD2 in flpIN CHO cells. _i_[Ca^2+^] quantified as a peak % of ionomycin following pramipexole application in DRD2 (orange, n=5), GHSR (green, n=5), and DRD2+GHSR (brown, n=5) transfected cells. (**B**) The presence of DRD2 does not change coupling of GHSR to calcium in flpIN CHO cells _i_[Ca^2+^] quantified as a peak % of ionomycin following ghrelin application in DRD2 (orange, n=5), GHSR (green, n=5), and DRD2 + GHSR (brown, n=5) transfected cells. (**C**) Quantification of EC_50_ and E_max_ of _i_[Ca^2+^] in DRD2 (orange) and DRD2+GHSR (brown) transfected cells following dopamine (main Fig **2D**), NOTE: of the 5n experiments curves could only be reliably fitted for 3n in DRD2 only cell line. (**D-G**) Quantification of EC_50_ and E_max_ of _i_[Ca^2+^] in DRD2 (orange) and DRD2+GHSR (brown) transfected cells following dopamine(**D,E**) or ghrelin (**F,G**) application, based on data in (**A** and **B**).. (**H,I**) The presence of GHSR does not alter DRD2 coupling to cAMP inhibition. Trace shows forskolin (FSK; 3 µM) mediated increase of cAMP (BRET/BRET max) which is reduced by pramipexole (100 nM) in cells transfected with DRD2+GHSR (brown) or with DRD2 (orange), mean - SD, n=5. (**I**) shows quantification using area under the curve (AUC) % of FSK at different concentrations of dopamine. Mean ± SD. (**J,K**) The presence of either receptor does not alter the cell surface expression of the other in an alternative model cell line. Cell surface expression of DRD2 (**J**) and GHSR (**K**) in single copy stably transfected flpIN HEK cell lines using flow cytometry and expression change (fold over control), DRD2 (orange), GHSR (green) and DRD2+GHSR (brown), representative of n=3. (**L**) GHSR expression is permissive for _i_[Ca^2+^] of DRD2 in an alternative model cell line (flpIN HEK cells). _i_[Ca^2+^] quantified as a AUC % of ionomycin following dopamine application in DRD2 (orange, n=4), GHSR (green, n=4), and DRD2+GHSR (brown, n=5) transfected cells. (**M**) DRD2 expression does not change _i_[Ca^2+^] of GHSR in an alternative model cell line (flpIN HEK cells). _i_[Ca^2+^] quantified as a AUC % of ionomycin following capromorelin application in DRD2 (orange, n=4), GHSR (green, n=4), and DRD2+GHSR (brown, n=5) transfected cells. All data mean ± SD * <0.05 student’s T-test.

**Fig. S3.**
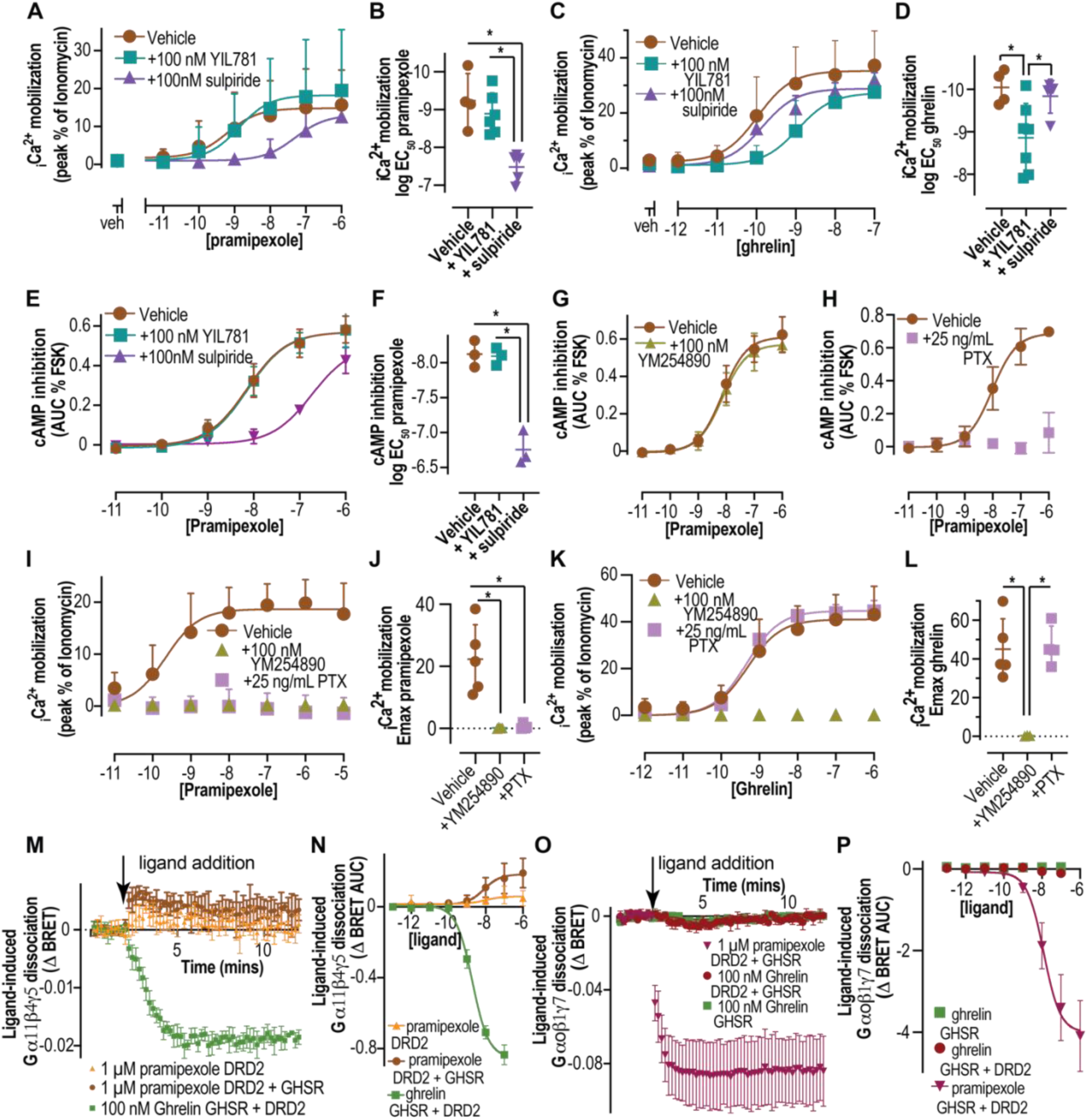
GHSR and DRD2 do not share a common signalling pathway in recombinant cell lines. (**A-D**) The effect of DRD2 preferring antagonist, sulpiride and GHSR antagonist, YIL781 on pramipexole (**A**) and ghrelin (**C**) dependent _i_[Ca^2+^] in cells co-expressing both receptors, shown as quantitated concentration-response curves of peak % of ionomycin response along with quantified EC_50_ (**B,D**) with one way ANOVA (n=5, with individual EC_50_ determinations in scatter plot). (**E, F**) The effect of DRD2 preferring antagonist, sulpiride and GHSR antagonist, YIL781 on pramipexole-dependent inhibition of cAMP production, concentration-response curves AUC % inhibition of FSK response (**E**) with quantified EC_50_ (**F**)(n=3, with individual EC_50_ determinations in scatter plot). (**G,H**) The effect of Ga_q/11_ (YM254890, **G**) or Ga_i/o_ (PTX, pertussis toxin, **H**) inhibition on dopamine-dependent inhibition of cAMP production, concentration-response curves AUC % inhibition of FSK response (n=3). (**I-L**) The effect of Ga_q/11_ (YM254890) or Ga_i/o_ (PTX) inhibition on pramipexole- (**I,J**) or ghrelin- (**K,L**) dependent _i_[Ca^2+^], as concentration-response curves peak % of ionomycin response (**I,K**) with quantified E_max_ (**J,L**)(n=5). (**M,N**) The presence of GHSR is not permissive for DRD2 coupling to Ga_11_/β4γ5 using TRUPATH G protein dissociation assay, (**M**) kinetic traces of BRET change in response to saturating ligand and (**N**) concentration response curves. (**O,P**) The presence of DRD2 is not permissive for GHSR coupling to Ga_o_/β1γ7 using a TRUPATH G protein dissociation assay, (**O**) kinetic traces of BRET change in response to saturating ligand and (**P**) concentration response curves. All data are mean ± SD or mean +SD; * p<0.05; one-way ANOVA or unpaired t-test, with estimation plot.

**Fig. S4.**
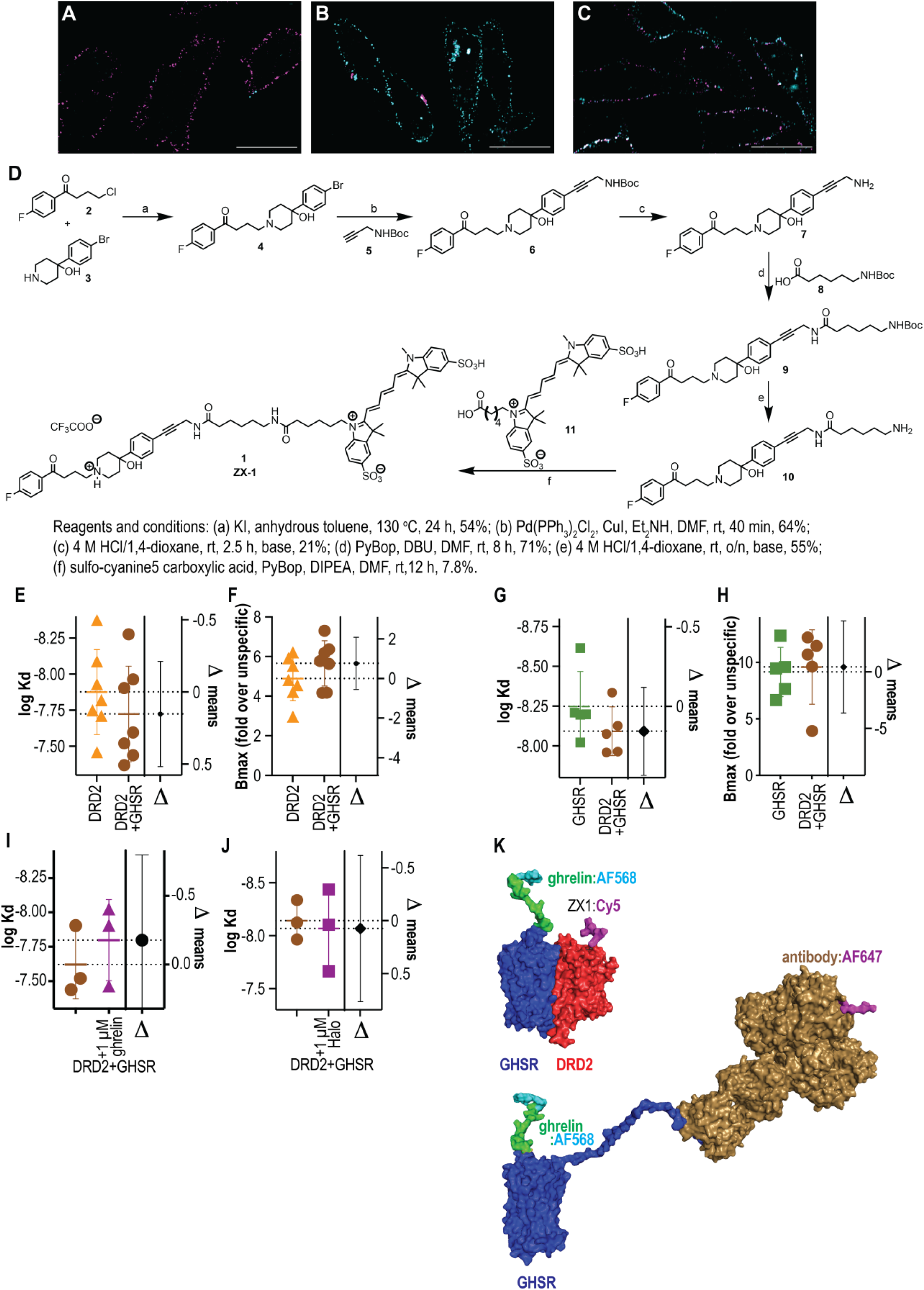
No detectable protein: protein interaction between GHSR and DRD2 in recombinant cell lines. (**A-C**) representative specificity controls for deconvolution microscopy (n=3) using direct conjugate antibodies directed toward n-terminal tags of DRD2 (pseudo-colored in magenta) and GHSR (pseudo-colored in cyan) with effective resolution, xy=160 nm, z=385 nm, scale bars are 20 µM. (**A**) GHSR only cells stained with both antibodies, (**B**) DRD only cells stained with both antibodies and (**C**) DRD2+GHSR cells stained with both antibodies. (**D**) Synthesis scheme (see methods) and structure for ZX-1 (haloperidol:Cy5). (**E,F**) estimation plots from individual saturation binding fits for Log Kd and Bmax values for ZX-1 binding to DRD2 (orange)and DRD2+GHSR (brown) cell lines (data from main Fig **3C**). (**G,H**) estimation plots from individual saturation binding fits for Log Kd and Bmax values for ghehlin:AF568 binding to GHSR (green) and DRD2+GHSR (brown) cell lines (data from main Fig **3D**). (**I**) estimation plots from individual saturation binding fits for Log Kd values for ZX-1 binding to DRD2+GHSR cell lines in the absence (brown) or presence (magenta) or 1µM ghrelin (data from main Fig **3E**). (**J**) estimation plots from individual saturation binding fits for Log Kd values for ghehlin:AF568 binding to DRD2+GHSR cell lines in the absence (brown) or presence (magenta) or 1µM haloperidol (Halo)(data from main Fig **3F**). (**K**) model of GHSR(blue)-DRD2(red) heterodimer based on TMIV-TMV interface with ghrelin modelled in based on solution structure of full-length peptide with AF568 appended to the end and ZX-1(black) extended from the haloperidol core from 6LUQ to the sluphoCy5(magenta) compared with a model of GHSR(blue) with ghrelin modelled in based on solution structure of full-length peptide with AF568 appended and n-terminus of GHSR as beta strand bound with a mouse IgG(gold) with AF647(magenta) appended to a surface exposed lysine in the Fc region to illustrate the differences in probable fluorophore proximity.

**Fig. S5.**
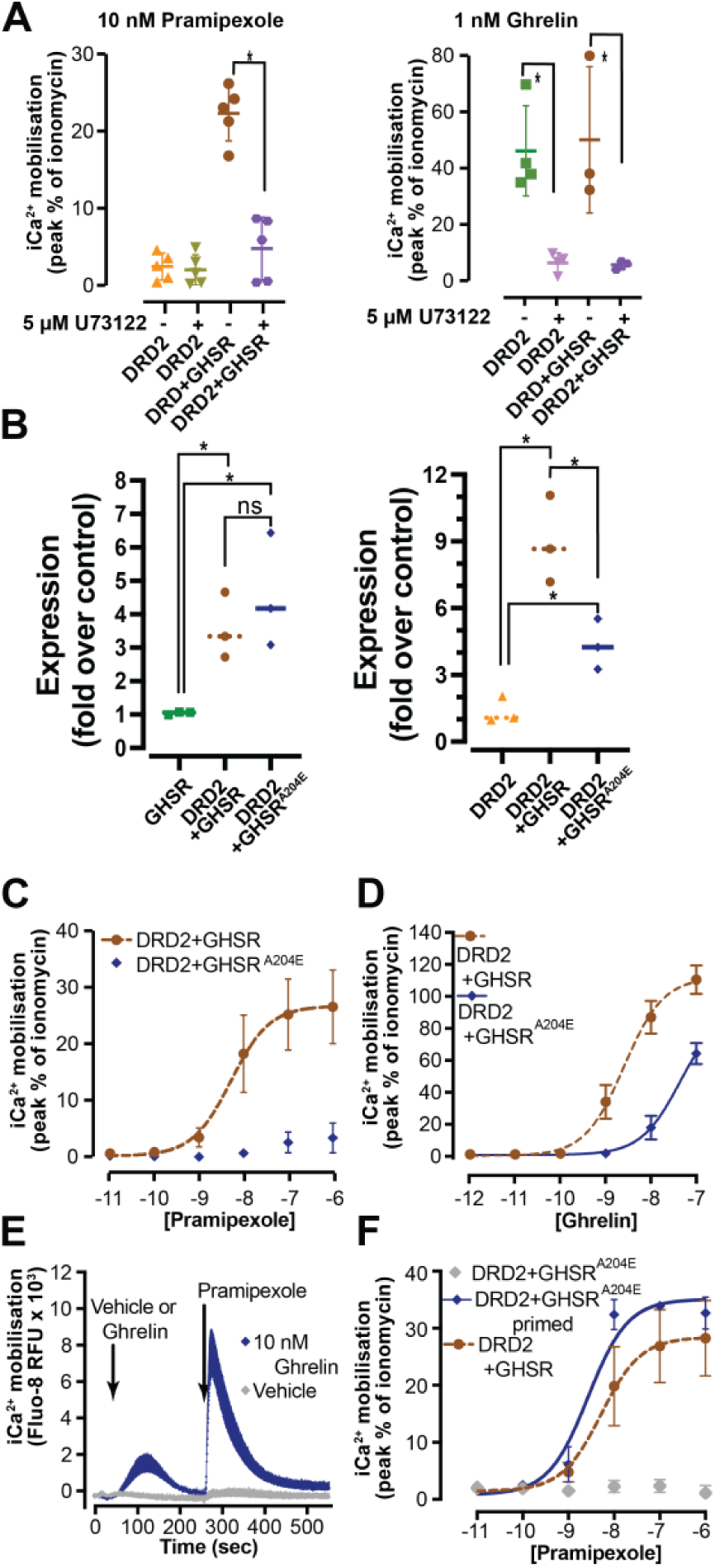
Requirement for PLC-β and ability of pre-stimulation to recover DRD2 coupling to calcium in transfected cells. (**A**) In transfected cells, co-addition of 5µM U73122 (PLCβ inhibitor) largely blocks the ability of pramipexole and ghrelin to evoke intracellular Ca^2+^ mobilization, shown as scatter plot of peak calcium (n=5). (**B**) Cell surface expression of DRD2 not altered by co-expression of GHSR^A204E^. Cell surface expression of DRD2 (left) and GHSR (right) in single copy stably transfected flpIN CHO cell lines using flow cytometry and pooled (fold over control), respectively, in DRD2 (orange, n=3), GHSR (green, n=3), DRD2+GHSR (brown, n=3) DRD2+GHSR^A204E^ (blue, n=3). (**C**) DRD2 fails to couple to Ca^2+^ in the presence of GHSR^A204E^ following pramipexole stimulation (**D**) and there is reduced GHSR calcium coupling in the presence of GHSR^A204E^. Mean ± SD. (n=3). (**E**) DRD2 Ca^2+^ coupling is rescued by pre-stimulation of GHSR^A204E^ with low concentration ghrelin. Representative trace showing pre-stimulation with ghrelin (10 nM; blue, n=3) eliciting a small Ca^2+^ response compared to vehicle (gray), followed by pramipexole leading to a sharp rise and substantial increase in Ca^2+^ release, compared to vehicle (gray). (**F**) Dose-response relationship of Ca^2+^ increase from pramipexole following pre-stimulation with ghrelin (10nM; blue, n=3).

